# Criticality Creates a Functional Platform for Network Transitions between Internal and External Processing Modes in the Human Brain

**DOI:** 10.1101/2020.12.25.424325

**Authors:** Minkyung Kim, Hyoungkyu Kim, Zirui Huang, George A. Mashour, Denis Jordan, Rüdiger Ilg, UnCheol Lee

## Abstract

Continuous switching between internal and external modes in the brain is a key process of constructing inner models of the outside world. However, how the brain continuously switches between two modes remains elusive. Here, we propose that a large synchronization fluctuation of the brain network emerging only near criticality (i.e., a balanced state between order and disorder) spontaneously creates temporal windows with distinct preferences for integrating internal information of the network and external stimuli. Using a computational model and empirical data analysis during alterations of consciousness in human, we present that synchronized and incoherent networks respectively bias toward internal and external information with specific network configurations. The network preferences are the most prominent in conscious states; however, they disrupt in altered states of consciousness. We suggest that criticality produces a functional platform of the brain’s capability for continuous switching between two modes, which is crucial for the emergence of consciousness.

## Introduction

Continuous switching between internal and external modes allows neural circuits to identify the contrast between information from two modes and reduce the mismatch between them (*1*, *2*). Continuous switching between two modes has been considered as a functional basis at a system level for constructing inner models in the brain, which supports perception, prediction, and action in the external world (*3*, *4*). However, it is unknown whether these transitions represent a necessary process to support higher-order brain functions, the origin of such modes in the brain, and the mechanism by which these modes transition. In our previous computational model study, we demonstrated that the brain’s responsiveness to external stimuli depends on the level of global brain network synchronization and this dependence only emerges near a critical state (*5*). In this study, we expanded the previous computational model study, suggesting that a large synchronization fluctuation emerging near a critical state may produce a functional platform, upon which functional brain networks fluctuate between two distinct modes: one of which conducive to the integration of internal information in the network and the other highly susceptible to external stimuli. Such distinct preferences for internal and external information of the brain originate from the general property of the network’s responsiveness with the synchronization fluctuation. We also analyzed both high-density electroencephalogram (EEG) and functional magnetic resonance imaging (fMRI) data of various states of consciousness (conscious, anesthetized, psychedelic, and pathological states) to investigate how the network’s preference for internal or external information in the time domain is associated with the altered states of consciousness.

Recent computational modeling and empirical studies suggest that consciousness occurs when brain dynamics are near criticality (i.e., poised at the border of multiple states) and that losing criticality (i.e., after a transition to one of the possible states) is related to altered states of consciousness (*6*–*10*). Critical dynamics confer biological advantages that may establish a functional foundation for the emergence of consciousness: an optimal balance between stability and instability, optimal computational capability, flexibility to adapt to a changing environment, and wide repertoires of brain states (*11*, *12*). In both biological and non-biological systems, a large global fluctuation is one of the most representative signal characteristics of criticality, with an increase in autocorrelation (*13*, *14*). In our previous brain network modeling study, we found that a large synchronization fluctuation near a critical state is associated with a highly variable brain sensitivity to external stimuli (*5*). Specifically, low and high levels of synchronization in the brain network respectively provide susceptible and refractory time windows to the stimuli. In addition, it has been suggested that the level of neural synchronization reflects the brain’s capability for information transmission and integration across the cortex (*15*, *16*). Based on these findings, we hypothesize that the functional brain network may be highly susceptible to external stimuli but less internally integrative at low levels of synchronization. Conversely, at high levels of synchronization, the brain network favors information integration within the network but is less susceptible to external stimuli. Thus, we hypothesized that the brain network at high and low levels of network synchronization possesses distinct preferences for, respectively, internal and external information, which may induce the internal and external modes of the brain in the time domain. Such distinct preferences may be the most significant in conscious states (i.e., near criticality) with the maximal difference between high and low levels of synchronization. In contrast, the preferences may be mitigated in altered states of consciousness (i.e., sub- or super-critical states) with reduced synchronization fluctuation.

To test the hypothesis, we used a large-scale human brain network model, modulating criticality and examining whether brain networks at high and low levels of synchronization near criticality have distinct preferences for internal information of the network and external stimuli, respectively. Symbolic mutual information (SMI) (*17*) and network susceptibility (*18*) were measured for each sub-second temporal window of simulated brain signals to evaluate quantitatively the brain network’s preference. The modeling results were tested in humans with high-density EEG during various states of consciousness; conscious wakefulness, anesthetized (with isoflurane and ketamine), psychedelic (with subanesthetic-dose of ketamine), and pathological conditions such as minimally conscious state and unresponsive wakefulness syndrome. We also examined functional brain network configurations at high and low levels of synchronization that systematically contribute to those preferences. In addition, we analyzed the fMRI data during wakefulness, propofol-induced unconscious state, and unresponsiveness wakefulness syndrome to examine whether the results observed in the model and EEG data are also consistent in the fMRI functional networks. We investigated whether the fMRI functional networks that predominate at high or low levels of synchronization are relevant to the well-known networks presumably associated with internal or external information processing. The schematic diagram of the study is presented in Fig. 1. The figures from fMRI data analysis were adapted from (*19*).

**Fig. 1.**
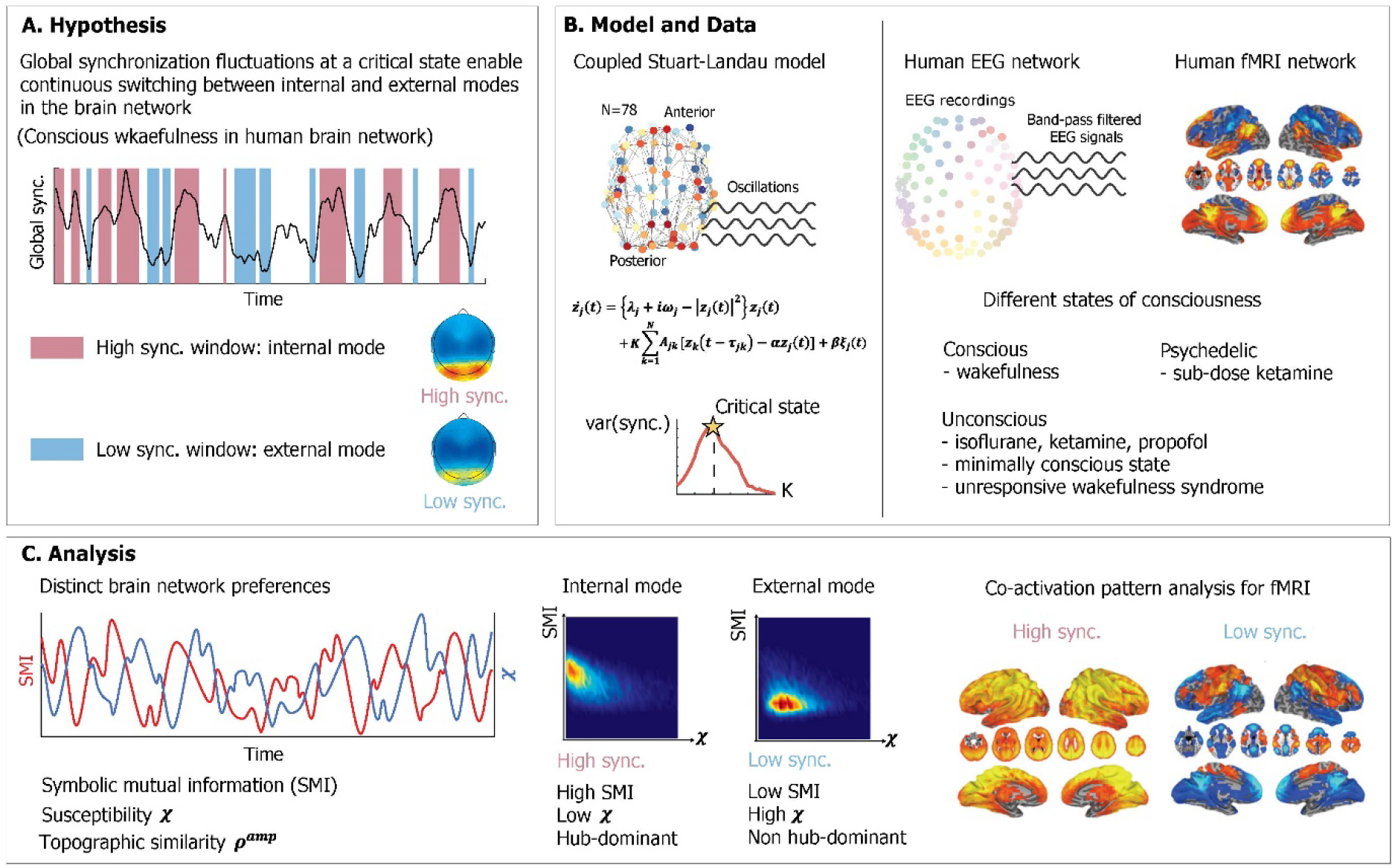
Schematic diagram of study design. We hypothesized that a large global synchronization fluctuation emerging near a critical state enables continuous switching between internal (;conducive to integration of internal information) and external (;highly susceptible to external stimuli) modes with distinct functional network configurations in a human brain network during conscious wakefulness. The study was composed of two parts, a mathematical modeling and an experimental analysis. We constructed a human brain network model, applying a coupled Stuart-Landau model to a human brain network structure consisting of 78 brain regions. The critical state was found with a maximum variance of the global synchronization. Human electroencephalogram (EEG) data were acquired in different states of consciousness including conscious states (CS), psychedelic state induced by sub-dose of ketamine, unconscious states (UCS) induced by isoflurane and ketamine, minimally conscious state (MCS) and unresponsive wakefulness syndrome (UWS). Functional magnetic resonance imaging (fMRI) data were acquired during CS, UCS induced by propofol and UWS. With the model and EEGs, we measured symbolic mutual information (SMI) and susceptibility *χ* in a sub-second time window to investigate the brain network’s preference for internal and external mode. The topographic similarity *ρ*^*amp*^ was measured to observe functional network configurations for each high and low synchronization window. The co-activation pattern analysis was performed for the fMRI data to obtain fMRI networks related to internal and external modes.

## Results

### Criticality produces distinct preferences for internal information and external stimuli in the brain network: a computational model study

We proposed that criticality in the brain network produces distinct preferences for internal information of the network and external stimuli, which may result from the general property of the network responsiveness with high and low levels of global network synchronization. By contrast, the preferences may vanish when the brain network is positioned far from criticality. To test this hypothesis, we used a large-scale human brain network model, simulating various brain network behaviors near and far from criticality.

The coupled Stuart-Landau model, which has successfully described the characteristics of EEG, magnetoencephalography (MEG), and fMRI in different states of consciousness, was used in this study (*9*, *20*–*23*). Spontaneous oscillations of the brain network were simulated with one hundred different initial frequency configurations and different sets of model parameters including the bifurcation parameter *λ*, coupling strength *K*, and diffusive coupling *α* (See “Materials and Methods” for details). Different parameter sets produce different brain states near and far from criticality. The level of network synchronization *R* was measured with a temporal average of the order parameter *r*(*t*). Fig. 2a presents the *R,* which is an average *r*(*t*) over 60 seconds, in the parameter space (λ, K). To capture the representative brain states near and far from criticality, we selected *λ* = −0.6 (dotted line in Fig. 2a) and the diffusive coupling of *K* = 0.5.

**Fig. 2.**
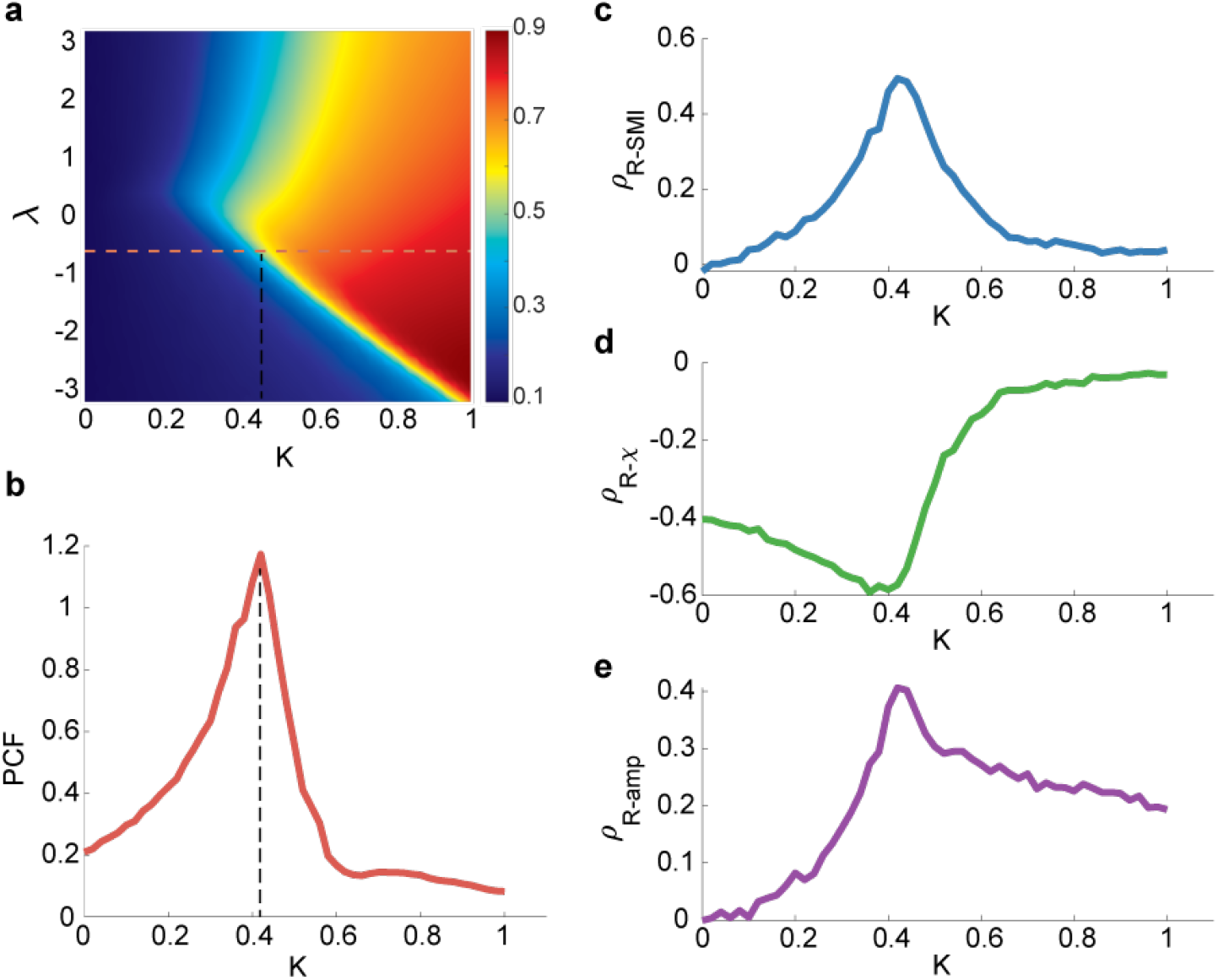
Relationships between network synchronization, symbolic mutual information, network susceptibility, and topographic similarity in the brain network model near and far from criticality. **a.** The relationship was investigated for a selected parameter set (the dotted line at *λ* = −0.6 and *K* = 0.42) in the model parameter space (*λ*: bifurcation parameter; *K*: coupling strength). The blue (red) color indicates a lower (higher) level of average network synchronization. **b.** Pair correlation function (PCF), a surrogate indicator of criticality, was maximal at the coupling strength *K* ≅ 0.42. Accordingly, a critical state is defined with the maximum PCF. Spearman correlations between network synchronization *R* and **c.** symbolic mutual information *SMI*, **d.** network susceptibility *χ*, and **e.** topographic similarity *ρ*^*amp*^ were calculated where *λ* is fixed at −0.6. The brain network model shows a maximum positive (negative) correlation between *R* and *SMI* (*R* and *χ*) near the critical state. The *ρ*^*amp*^ also shows a maximum positive correlation with *R* near the critical state.

Fig.2b presents a change of pair correlation function (PCF), which measures a temporal variance of network synchronization as a surrogate indicator of criticality (*18*), along with increasing *K*. The critical state was defined with the maximal PCF, indicating the largest synchronization fluctuation at criticality. Here, we defined the brain state at *K* ≅ 0.42 as one of the critical states in the model. The other critical states with a different parameter set present qualitatively similar features (Fig. S1-S3).

The brain network’s preferences for integrating internal information of the network and external stimuli were respectively evaluated with symbolic mutual information *SMI* (*17*) and network susceptibility *χ* (*18*). *SMI* quantifies the amount of shared information and *χ* quantifies network susceptibility to external stimuli. Fig. 2c and Fig. 2d show that both measures have maximal Spearman correlation coefficients with *R* (*ρ*_*R-SMI*_ = 0.50 *K* = 0.42 and *ρ*_*R-χ*_ = −0.59 at *K* = 0.38, respectively) near the critical state. In other words, a brain network with a larger *R* has a larger *SMI* and smaller *χ* (preference for integrating internal information of the network), whereas a brain network with a smaller *R* has a smaller *SMI* and larger *χ* (preference for external stimuli). Here, *R*, *SMI*, and *χ* were calculated within each 250-ms temporal window with 50-ms overlap, and the correlations among them were calculated across the temporal windows (See “Materials and Method” for the details). Note that the sub-second temporal scale is a suitable time resolution for the information integration processing in a large-scale network level (*24*).

We also examined the functional network configurations of high and low *Rs* to investigate the functional network basis for two different preferences. We first calculated topographic similarity *ρ*^*amp*^, which is defined as a Spearman correlation between node degrees of the anatomical brain network and average amplitudes of the simulated brain activities. In this analysis, a larger *ρ*^*amp*^ implies that hub nodes are more dominant in network dynamics along with higher amplitudes. Then we calculated the Spearman correlation coefficient between *ρ*^*amp*^ and *R*. Fig. 2e shows that the correlation between *R* and *ρ*^*amp*^ is also maximal (*ρ*_*R-amp*_ = 0.41) near the critical state, suggesting that a highly synchronized brain network is more likely to have a hub-dominant functional network configuration that is optimal for integrating internal information of the brain network. The results are consistent with different diffusive coupling parameters, *α* = 0 and *α* = 1 (Fig. S2 and Fig. S3).

### The preferences for internal information and external stimuli of the brain network are prominent near criticality in a sub-second time window: a computational model study

To test whether the preferences of the brain network are significantly different in the time domain, we classified the simulated brain signals into low and high *R* windows and investigated *SMI* and *χ* values for each temporal window (250-ms). Fig. 3a shows a temporal evolution of *R* during 60 seconds in the brain network model near a critical state. The low *R* and high *R* windows were defined with thresholds *R* < 0.3 and *R* > 0.5, respectively (blue: low *R* and red: high *R* in Fig. 3a). The high *R* windows are characterized by large *SMI* and small *χ*, whereas the low *R* windows have relatively small *SMI* and large *χ* (***p<0.001 for SMI; ***p<0.001 for *χ*, Wilcoxon rank-sum test, in Fig. 3b and 3c). The results show that the brain states of these high and low *R* windows define preferences for internal information vs. external responsiveness in the network. We also showed that high *R* windows have a large positive *ρ*^*amp*^ ; by contrast, low *R* windows have negative *ρ*^*amp*^ (***p<0.001, Wilcoxon rank-sum test, Fig. 3d). The *ρ*^*amp*^ value itself was variable in different parameter sets, but the positive correlation between *R* and *ρ*^*amp*^ near criticality was consistent (Fig. S1-S3). Our results suggest that the fluctuation of network synchronization near criticality naturally produces a continuous switching between internal and external modes in the brain network with different network configurations. In Fig.3e, the joint histograms of SMI and *χ* (see “Materials and Methods” for more details) clearly show the distinct preferences of high (right) and low (left) *R* windows for internal information of the network and external stimuli (i.e., larger *SMI* and smaller *χ* for high *R* windows; smaller *SMI* and larger *χ* for low *R* windows), respectively.

**Fig. 3.**
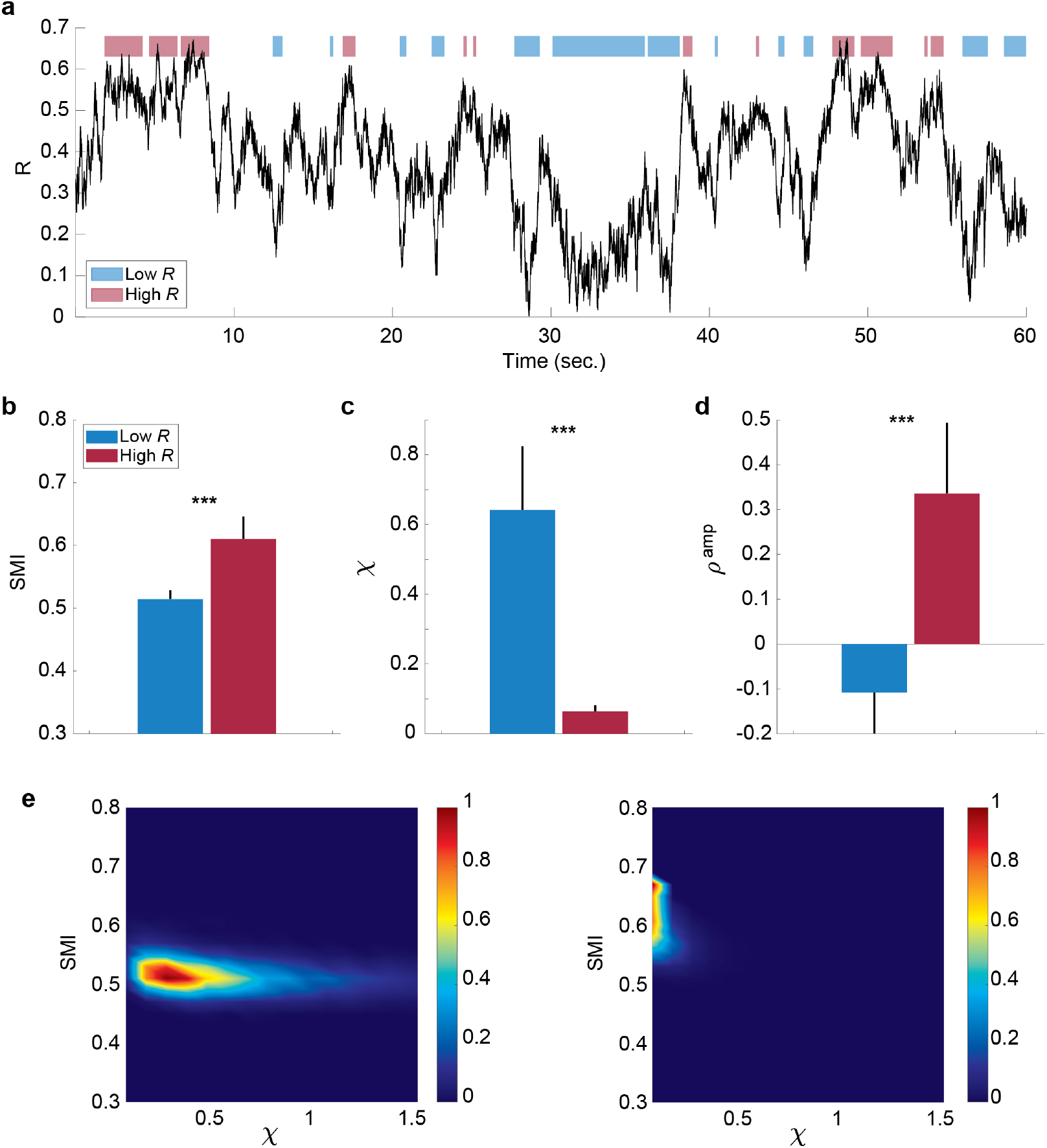
The preferences of the brain network for internal information integration and external responsiveness are prominent in sub-second time windows (250-ms). **a.** An example of a network synchronization fluctuation near a critical state. Red and blue bars indicate temporal windows classified into low and high *R*. The *SMI*, *χ*, and *ρ*^*amp*^ were calculated for each window. Comparisons of **b.** *SMI*, **c.** *χ*, and **d.** *ρ*^*amp*^ between temporal windows of low *R* (blue) and high *R* (red) are presented. The black lines of the colored bars indicate from 25% to 75% quantiles. A Wilcoxon rank-sum test was performed (***p<0.001). **e.** Joint histograms of the low (left) and high (right) *R* windows. The low *R* window is characterized by relatively low *SMI* and high *χ*, while the high *R* window is characterized by relatively high *SMI* and low *χ*.

### The preferences of the brain network for internal information and external stimuli are prominent in the conscious state: an EEG study

To test the modeling results empirically, we analyzed EEG signals of conscious eye closed-resting states in humans (n=24). Computational model and empirical studies suggested that theta and alpha-band (4-12 Hz) oscillations are globally networked in the cortex and spatiotemporally organize neural processing with traveling waves across the human cortex (*25*, *26*). Thus, here we focused on the EEG signals of 4-12 Hz which is suitable to test the characteristic behavior of the brain network near criticality. We also observed that the results of this frequency band (4-12 Hz) are most consistent with the model predictions (Fig. S4 and S5). As in the model data analysis, we calculated *R*, *SMI*, *χ*, and *ρ*^*amp*^ for each temporal window of 250-ms. Fig. 4a presents a temporal evolution of *R* for 60 seconds. We used thresholds 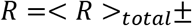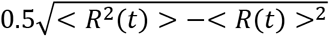, where < *R* >_*total*_ is an average of *R* across all subjects and all states of EEGs (See “Materials and Method” for more details), to classify the temporal windows into high *R* and low *R* windows (blue and red blocks in Fig. 4a). Note that we used different thresholds for the source signals (model) and sensor signals (EEG) due to the differences in their signal variability. The results of the EEG analysis were consistent with the results of the computational model and show l arge *SMI* and small *χ* for high R windows and small *SMI* and large *χ* for low *R* windows (***p<0.001 for *SMI*; **p<0.01 for *χ*, Wilcoxon rank-sum test, Fig. 4b and Fig. 4c). The high *R* windows also presented a large positive *ρ*^*amp*^ (**p<0.01, Wilcoxon rank-sum test, in Fig.4d). For *ρ*^*amp*^, we used average degrees of weighted phase lag index (wPLI) networks and amplitudes of EEG signals from 96 channels, assuming that the wPLI network averaged over a long time may resemble its underlying structural network (*22*). Similar to the brain network model near criticality, the brain network during conscious states demonstrates the distinct preferences of high and low *R* windows for internal information and external stimuli as well as the posterior hub-dominant network configuration in high *R* windows (Fig. 4e). Furthermore, joint histograms of *SMI* and *χ* clearly visualizes the distinctive preferences of high (right) and low (left) *R* windows in conscious states (Fig. 4f). In sum, the brain network constructed from EEGs during conscious states shows the typical network properties of high and low *R* windows in terms of *SMI* and *χ*, which is predicted by the model study, and temporally yields continuous switching between preferences for internal information and external stimuli.

**Fig. 4.**
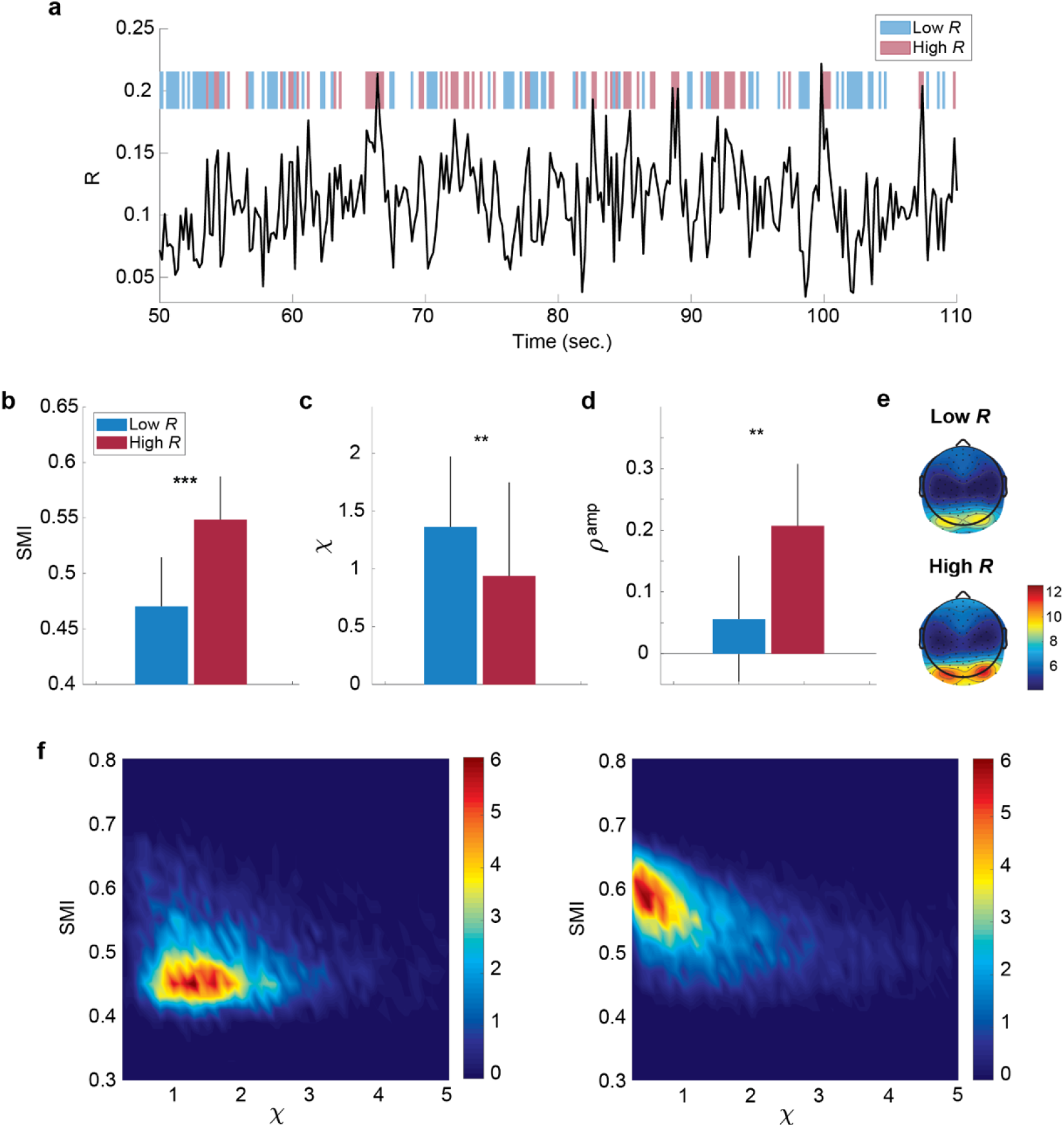
EEG functional brain networks in conscious state present distinct preferences for internal information integration and external responsiveness in the time domain. **a.** An example of a network synchronization fluctuation during a conscious state. Red and blue bars respectively indicate temporal windows classified into low and high *R*. The *SMI*, *χ*, and *ρ*^*amp*^ were calculated for each 250-ms window. Comparisons of **b.** *SMI*, **c.** *χ*, and **d.** *ρ*^*amp*^ between low *R* (blue) and high *R* (red) windows are presented. The black lines of the colored bars indicate from 25% to 75% quantiles. A Wilcoxon rank-sum test was performed (***p<0.001 and **p<0.01). **e.** Functional brain network configurations for low and high *R* windows. As expected in the model study, the amplitudes of the posterior area (hub regions) in high *R* windows are larger than those in low *R* windows. **f.** Joint histograms of the temporal window of low (left) and high (right) *R*. The low *R* windows are characterized by relatively low *SMI* and high *χ*, while the high *R* windows are characterized by relatively high *SMI* and low *χ*. The empirical data are consistent with the model prediction.

### The preferences of the brain network for internal information and external stimuli vanishes in altered states of consciousness: an EEG study

Our modeling results predict that the distinct preferences of high and low *R* windows for internal information and external stimuli vanish when the brain network deviates from criticality, with reduced synchronization fluctuation (Fig. 2). To test this model prediction, we analyzed EEGs from conscious eyes-closed resting states in humans (CS, n=24), isoflurane-induced unconsciousness (ISO, n=9), ketamine-induced unconsciousness (KET, n=15), subanesthetic ketamine-induced psychedelic state (PSY, n=15), and disorders of consciousness such as minimally conscious states (MCS, n=16) and unresponsive wakefulness syndrome (UWS, n=9). We first calculated the PCF to evaluate the synchronization fluctuation and demonstrated that the PCF significantly decreases in all altered states of consciousness (p<0.001, Kruskal-Wallis test, Fig. 5a). The PCFs of the five altered states of consciousness are significantly different from one another (p<0.001, multiple-comparison test with Tukey-Kramer method) except for the ISO and PSY (p=0.59) pair and the KET and MCS pair (p=0.08).

**Fig. 5.**
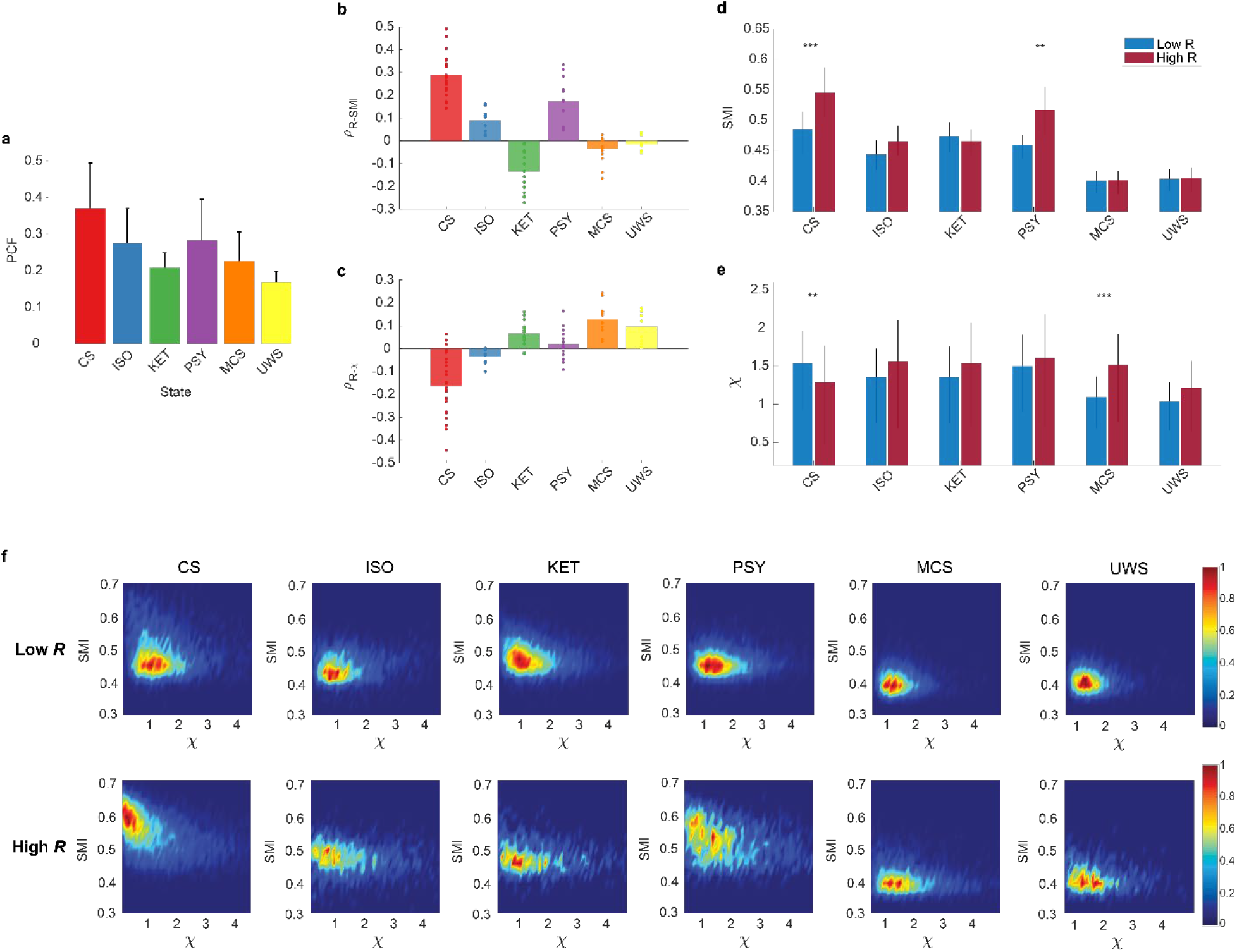
The preferences of the brain network for internal information integration in the network and external responsiveness are significantly disrupted in altered states of consciousness (CS: conscious state, ISO: isoflurane-induced unconsciousness, KET: ketamine-induced unconsciousness, PSY: subanesthetic ketamine-induced psychedelic state, MCS: minimally conscious state, and UWS: unresponsive wakefulness syndrome). **a.** PCF in different states of consciousness (Mean±SD). The CS shows the largest PCF as expected in the brain network model. PCFs are significantly different among the states except for the ISO and PSY pair and the KET and MCS pair (p<0.001, Kruskal-Wallis test with a multiple-comparison test using Tukey-Kramer method). Spearman correlations between **b.** *R* and *SMI* and **c.** *R* and *χ* in different states of consciousness. Dots indicate the correlation values of subjects. As expected in the model study, CS shows a maximum positive (negative) correlation between *R* and *SMI* (*R* and *χ*), and these relationships are disrupted in the altered states of consciousness. **d.** *SMI* and *χ* for low and high *R* windows in different states of consciousness. Only CS shows significant differences between high and low *R* windows in both *SMI* and *χ* (***p<0.001 and **p<0.005; Wilcoxon rank-sum test). **f.** Joint histograms of different states of consciousness. The distinct preferences of low and high *R* windows disrupt in all the altered states of consciousness.

We also calculated the *R*, *SMI*, and *χ* for each temporal window and calculated the correlations between them. With reduced PCFs, as expected, all of the altered states of consciousness lost the correlations between *R* and *SMI* and between *R* and *χ*, except for *R* and *SMI* of PSY (Fig. 5b and 5c). While investigating several other frequency bands (1-4 Hz, 1-20 Hz, 8-12 Hz, 13-30 Hz, and 30-42 Hz), we found that the frequency band of 8-12 Hz is also consistent with the results from 4-12 Hz (Fig. S4 and S5). Furthermore, we also classified the temporal windows of all states into high and low *R* windows with the same thresholds of CS. The distinct preferences between internal information and external stimuli observed in CS (large *SMI*/small *χ* for high *R* and small *SMI*/large *χ* for low *R* window) faded away in the altered states of consciousness (*SMI*; CS: ***p<0.001, ISO: p=0.03, KET: p=0.48, PSY: **p<0.005, MCS: p=0.90, and UWS: p=1, *χ*; CS: **p<0.005, ISO: p=1, KET: p=0.06, PSY: p=0.10, MCS: ***p<0.001, UWS: p=0.14, Wilcoxon rank-sum test) (Fig. 5d and Fig. 5e). We also found that the *SMI* of high *R* windows in CS is significantly reduced in ISO, KET, PSY, MCS, and UWS (p<0.001; CS vs. all other states, one-way ANOVA with multi-comparison test with Tuckey-Kramer method). By contrast, the *χ* of low *R* windows of ISO, KET, MCS, and UWS was significantly decreased or maintained compared to that of CS. This result implies that the brain capability for internal information integration might play a more prominent role in altered states of consciousness than network susceptibility to external stimuli. In Fig. 5f, joint histograms of *SMI* and *χ* visualize that only CS shows distinctive preferences of low and high *R* windows for internal information of the network and external stimuli. Functional network configurations for low and high *R* windows were also investigated in altered states of consciousness and presented in Supplementary materials (Fig. S6). Different from the CS, ISO shows anterior dominant network configuration for the high *R* windows and KET shows no difference between low and high *R* windows. As would be predicted, the network configuration for the high *R* windows in PSY is similar to the one during CS. Even though the R values are similar to each other in high *R* windows across the states, the posterior hub-dominant network configurations during CS play a pivotal role in brain’s distinct capability for internal information integration (Fig. S7).

### Dominant fMRI co-activation patterns at high and low levels of synchronization in conscious and unconscious states

Finally, we tested whether the preference of high and low *R* windows can be confirmed using a different brain imaging modality such as fMRI BOLD signals, investigating an association with well-known fMRI networks that are activated while the brain carries out tasks requiring internal and external information processing. We predicted that the fMRI networks relevant to internal or external information processing may be preferentially activated in high or low R windows and such preferences may change in altered states of consciousness.

To determine which fMRI networks are dominant in high and low *Rs*, we first classified human fMRI BOLD signals from conscious states (CS, n=42), propofol-induced unconscious states (Prop, n=14), and UWS (n=13) into low and high *R* windows with the thresholds 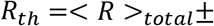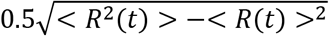, where < *R* >_*total*_ is an average network synchronization across all subjects and all states. We then investigated the occurrence rates of co-activation patterns (CAPs) for the low and high *R* windows, comparing them with the occurrence rates of a null CAP time-series generated by 20,000 permutations of BOLD signals (Fig. 6a). The CAPs were defined as 8 typical patterns of BOLD co-activation across voxels such as default mode network (DMN+), dorsal attentional network (DAT+), frontoparietal network (FPN+), sensory and motor network (SMN+), visual network (VIS+), ventral attention network (VAT+), and a global network of activation and deactivation (GN+ and GN-). More detailed explanation of the CAP analysis is in “Materials and Methods” and Supplementary materials Fig. S8. According to our hypothesis, the low *R* windows should be characterized by networks that are sensitive to external stimuli, whereas the high *R* windows should be more relevant to networks that are insensitive to external stimuli. Our results show that in CS the low *R* windows are dominated by DMN+, DAT+, and FPN+, which are sensitive to external stimuli (*27*), and the high *R* windows are dominated by GN+ and GN-, which are known as arousal networks that are insensitive to external stimuli (Fig. 6b) (*28*–*30*). The dominant CAPs in CS were replaced with other CAPs in Prop and UWS, mostly deactivating DMN+. Thus, the distinct preferences of high and low *Rs* for internal information of the brain network and external stimuli observed near criticality in the conscious state can be confirmed with fMRI BOLD signals. The disrupted preferences were also observed universally in the brain network that distant from criticality and the EEG and fMRI data in altered states of consciousness.

**Fig. 6.**
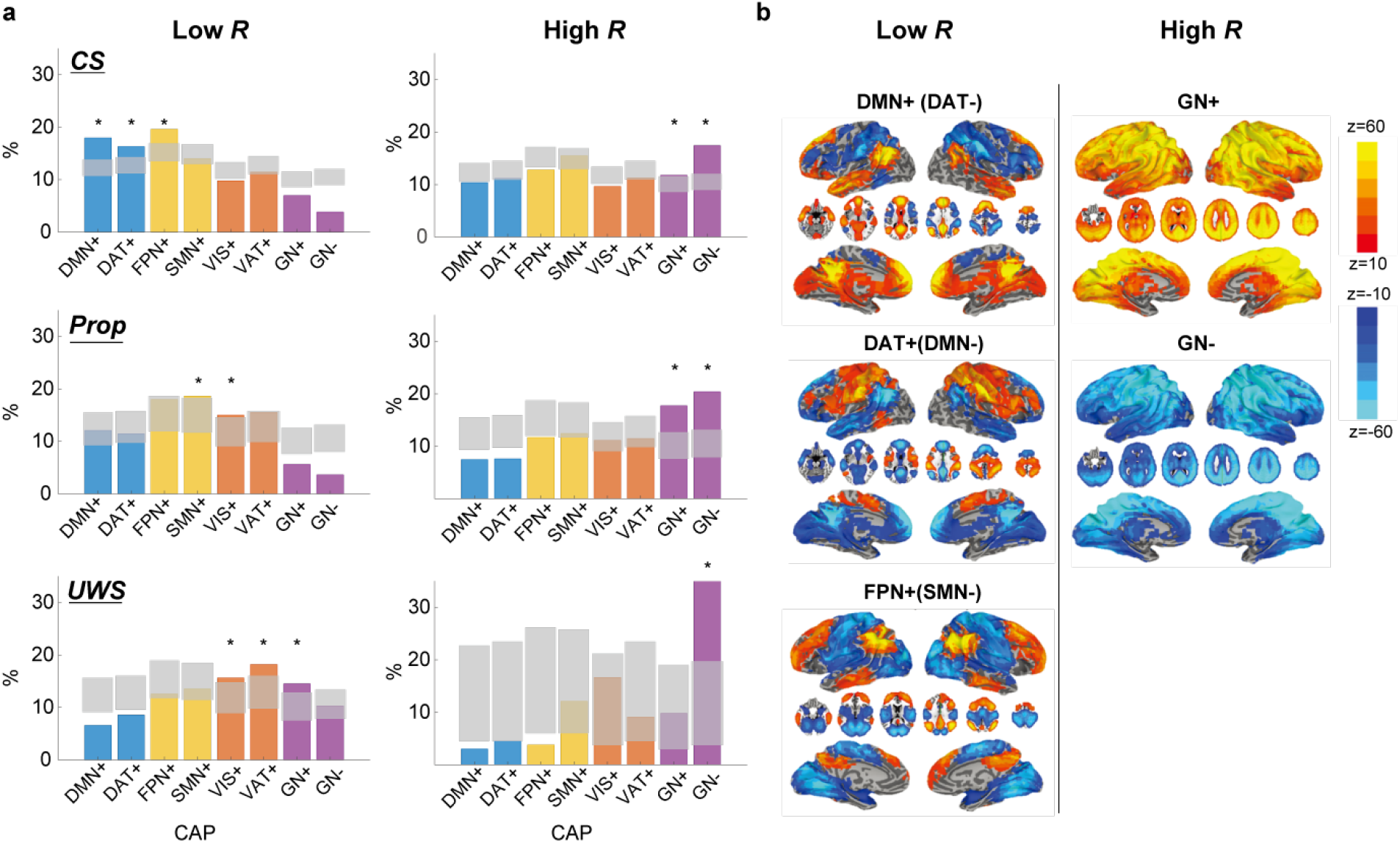
Dominant fMRI co-activation patterns (CAPs) for low and high *R* windows in conscious states (CS), propofol-induced unconsciousness (Prop), and unresponsive wakefulness syndrome (UWS). **a.** Occupancy percentages of CAPs in CS (1^st^ row), Prop (2^nd^ row), and UWS (3^rd^ row) for low *R* (left) and high *R* (right) windows. Gray areas indicate quantiles ranging from 0.5% to 99.5% from the distributions of 20000 permutation sets. Significantly dominant CAPs are marked as ‘*’ in the figure. The CAPs consist of eight different functional networks such as a default-mode network (DMN+), dorsal attention network (DAT+), frontoparietal network (FPN+), sensory and motor network (SMN+), visual network (VIS+), ventral attention network (VAT+), and a global network of activation and deactivation (GN+ and GN-). Functional networks including DMN+(DAT-), DAT+(DMN-), and FPN+(SMN-) are dominant in low *R* windows, while functional networks including GN+ and GN- are dominant in high *R* windows in CS. **b.** Spatial maps of CAPs for low and high *R* windows in CS. Spatial maps of other CAPs are presented in the supplementary materials (Fig. S8).

## Discussion

With a computational model, we demonstrated that when the brain network resides near a critical state—a balanced state between incoherent and synchronized connections (modulated by the coupling strength)—the global synchronization *R* correlates with the amount of shared information (*SMI*) within the network and the network susceptibility (*χ*) to external stimuli. The highly synchronized brain network is characterized by a large *SMI* and small *χ* (preference for internal information), while the incoherent brain network shows a small *SMI* and large *χ* (preference for external stimuli). However, the correlations between *R*, *SMI*, and *χ* were diminished when the brain network deviates from criticality. The modeling study also showed that a highly synchronized brain network displays predominant hub activities, which provides a network condition favorable for internal information integration. The modeling results were tested with the empirical data analyses across diverse states of consciousness. First, the brain network during baseline consciousness exhibited the largest synchronization fluctuation (i.e., the maximal PCF) compared to the altered states of consciousness (ISO, KET, PSY, MCS, and UWS), which implies criticality of the conscious brain. Second, the functional brain networks constructed from EEG during CS showed temporally variable preferences between internal information of the network and external stimuli, presenting a large *SMI*/small *χ* at high *R* and a small *SMI*/large *χ* at low *R*. These preferences were significantly diminished in the altered states of consciousness. Additionally, the dwell-time of the hub-dominant network configurations at high *R* windows were organized in a scale-free manner. The power-law distribution of the dwell-time changed into a random organization (i.e., exponential distribution) in unconscious states (Fig. S8). Finally, for the fMRI signals, the most frequently observed CAPs at high (low) *R* corresponded to internally integrative (highly susceptible) brain states.

In sum, distinct preferences for internal information of the network and external stimuli spontaneously emerge when the brain network is positioned near criticality and in conscious states. The continuous fluctuation between these two preferences may be a functional basis for constructing inner models of the outside world. When deviated from criticality and in altered states of consciousness, the brain loses such distinct preferences with significantly diminished synchronization fluctuation, which disrupts the capability for integrating internal information or receiving external information in the time domain.

### A novel role of criticality in conscious brain function

Biological systems can obtain many functional benefits from operating near a critical point of a phase transition. Criticality in biological systems, conjectured to emerge as the result of adaptive and evolutionary processes, produces an optimal balance between stability and instability, optimal computational capability, large dynamical repertoires, and greater sensitivity to stimuli (*6*–*9*, *11*, *12*). The brain near a critical state should display characteristic features that are empirically measurable: maximal sensitivity, large spatiotemporal correlations, and large variances in synchronization. Maximal sensitivity is an important property for the sensory systems in the brain, such as the olfactory, visual, and auditory systems, optimizing responses to environmental cues (*31*–*35*). A large spatiotemporal correlation is crucial for the brain to induce coordinated neural activities across space and time (*7*), which is a useful mechanism for the generation of long-lasting and slow-decaying memories at multiple timescales (*36*). A large variance of synchronization induces large statistical complexity and large repertoires in the brain, which is due to the maximal variety of attractors and metastability (*8*, *36*, *37*) in corresponding state space. It facilitates the spontaneous generation of complex patterns required for optimizing the brain’s capability for storing and processing information, enabling the brain to constantly traverse different network configurations, which is associated with cognitive flexibility. Based on our findings in the model study (Fig. 2 and Fig. 3) and the empirical data analysis (Fig. 4), here we propose a novel role of criticality for brain function. Near criticality, the large synchronization fluctuation in the brain network produces distinct preferences for internal information of the network and external stimuli, which enables the brain network to integrate internal information or be sensitive to external information in the time domain. Furthermore, the temporally variable network preferences might provide a functional platform for continuous switching between internal and external information modes in the brain, which is essential for constructing inner models of the outside world through the recursive learning process (*3*) as well as for the emergence of consciousness. In addition, as a network becomes distant from a critical state along with the reduced synchronization fluctuation, we can expect that such distinct preferences vanish, accompanying with the altered state of consciousness. Our model simulation and empirical data analyses explicitly demonstrate that the variance of synchronization (PCF) is maximal at a critical state (Fig. 2b) and in conscious states (Fig. 5a). By contrast, the variance is significantly diminished in altered states of consciousness such as general anesthesia, psychedelic experiences, and pathological disorders of consciousness (Fig. 5a).

### The brain network preferences vanish in altered states of consciousness

The brain networks in the altered states of consciousness no longer possessed distinct preferences for internal information of the network and external stimuli. Then, how do the brain networks in the altered states of consciousness lose such preferences? Is there a common network feature across the altered states of consciousness, and a distinctive network feature that characterizes each state?

The decreased *SMI*, especially in high *R* windows, is a common feature across altered states of consciousness regardless of the types of anesthetics and traumatic injuries. However, network susceptibility *χ* was relatively not changed across the states. These results imply that the brain network preferentially loses its capability to integrate internal information, while its susceptibility to external stimuli remains relatively intact. In other words, the brain network may be able to receive external stimuli but it cannot be globally integrated during the altered states of consciousness, which may be specifically related to functional deafferentation and disconnection for the external world due to the isolation of thalamocortical network (*38*). In addition, alterations of consciousness accompanied the disruption of posterior hub-dominant network configuration (Fig. S6), which impairs the functional role of hubs for integrating and transmitting information within the hierarchical brain network (*39*, *40*). This hub disruption also causes the selective inhibition of top-down processes through preferentially impeding information flow from hubs to peripherals in the brain network (*39*, *41*, *42*). Many anesthesia experiments have demonstrated that anesthetics selectively inhibit higher-order information integration in top-down processes while preserving the bottom-up and primary sensory processes, highlighting the importance of top-down processes for the emergence of consciousness (*43*, *44*). Our model and empirical data analyses suggest that such preferential inhibition of the internal information integration in the brain network, specifically, via disrupting the hub-dominant network configuration in high *R* windows, which is likely associated with high-order information integration, is a typical phenomenon that can occur when a complex network deviates from criticality. If the network moves to states far from criticality (super- or sub-critical state), it will preferentially hamper the normal functions of hubs that have dense connections.

The altered states of consciousness presented distinctively impaired network properties, *SMI* and *χ*, in high and low *R* windows, which can differentiate all the states. Interestingly, the psychedelic and minimally conscious states showed opposite network properties to one another (Fig. 5d). The psychedelic state preserved the capability for integrating internal information of the network (i.e., larger *SMI* in high *R* window), however, the discrimination of network susceptibility between high and low *R* windows vanished. The low *R* windows no longer retained the bias toward external stimuli. In contrast, the MCS lost the capability for integrating internal information (i.e., small *SMI* in high *R* window), but instead presented larger network susceptibility in high *R* windows, which is the opposite compared to waking consciousness (Fig. 5e). In other words, the brain network of MCS disrupted the capability for internal information integration with the abnormal relationship between network synchronization and susceptibility. Conversely, the brain network of psychedelic state still functions for internal information integration, however, it does not respond to external stimuli properly, which suggests an impaired network preference for external stimuli. Finally, we found that the dwell-time for the hub-dominant network configuration, follows a power-law distribution in the conscious state but follows an exponential distribution in during general anesthesia with either isoflurane or ketamine (at anesthetic levels) (Fig. S9). The results imply that the hub-dominant network configurations are temporally organized in a scale-free manner, which enables the brain to process higher-order information integration at various time scales. However, the altered states of consciousness significantly restrain potential neuronal repertoires in the brain network not only in the spatial domain but also in the time domain.

### Dominant fMRI co-activation patterns at high and low levels of synchronization

Dominant fMRI co-activation patterns for incoherent and synchronized windows in conscious states were presumably associated with the brain networks that are sensitive and insensitive to external stimuli, respectively. We showed that in conscious states the co-activation patterns at incoherent windows were dominated by DMN+, DAT+, and FPN+. These networks have complex interactions and support higher-order cognitive functions. For example, the DMN engages in a variety of processes such as autobiographical memory, imagination, and self-referencing (*45*). The DAT mediates cognitive processes such as goal-driven attention, inhibition, and top-down guided voluntary control (*46*, *47*). The FPN+ is known to flexibly alter its functional connections dynamically according to current task demands (*48*), and has a strong association with working memory (*49*). Furthermore, these three networks are at a high position of a representational hierarchy, relatively far from the sensory and motor systems in terms of both functional connectivity and anatomical distance (*50*). Such a hierarchical disposition is thought to allow those networks to process transmodal information in a way that is unconstrained by immediate external stimuli. We suggest that these spatially segregated co-activation patterns (DMN+, DAT+, and FPN+) may be associated with high sensitivity to stimuli. As supported by a previous study by Sadaghiani et al. (*27*), pre-stimulus brain states with higher modularity (i.e., higher spatial segregation and lower global integration) bias toward detecting external stimuli, whereas pre-stimulus brain states with lower modularity (i.e., higher spatial homogeneity and higher global integrity) bias toward misses.

In contrast, we found that the highly synchronized states in conscious states are dominated by GN+ and GN-. The two co-activation patterns are correlated with global EEG synchronization in the alpha frequency band (*28*), and associated with arousal fluctuations regulated by subcortical-cortical connectivity (*28*–*30*). Particularly, the GN+ has been suggested to be specific to lapses in alertness associated with a transition to a state of lower arousal (*28*), rendering the neural system less sensitive to external stimuli. Taken together, our results suggest that the distinct preferences of the brain network for internal information and external stimuli in low and high network synchronizations, regardless of imaging modalities, are consistent network properties near critical states.

### Potential mechanisms for the emergence of distinct preferences in the brain network

Recent studies of brain network characteristics near criticality have provided some evidence of the potential mechanism for the emergence of continuous switching between internal and external modes. One of the potential mechanisms is a global fluctuation in neural gain mediated by ascending neuromodulatory nuclei such as pontine locus coeruleus (*51*, *52*). A series of studies has suggested that the modulation of neural gain with the dynamic changes in noradrenaline results in a large fluctuation between network integration and segregation in the brain (*51*, *53*). When the neural gain dynamics (presynaptic afferent input) of the locus coeruleus reside near criticality, a small fluctuation produces a sharp transition between network integration and segregation, which respectively accompanies optimized information transfer and optimized information storage in the brain (*53*). Network integration and segregation were also associated with an increased spatial correlation (correlated with elevated information transfer) and increased autocorrelation times (correlated with increased information storage) with correspondence to high and low levels of phase synchronization (*51*, *53*).

A previous computational model study from our research team found that the brain network’s responsiveness significantly depends on the level of network synchronization as well as the distance from criticality when a stimulus is applied (*5*). Based on these results, we suggested that a potential mechanism for the relationship between brain network synchronization and responsiveness is phase response curve (PRC) in physics, which is a general property of networked oscillators ubiquitously observed in physical and biological systems. The PRC describes the way that a system with a collective periodic behavior (for instance, circadian rhythms, cardiac rhythms, and spiking neurons) responds to external stimuli (*54*, *55*). The response of an oscillating system is measured by the phase shift from the original phase and the phase shift (advancing or delaying the original phase) is an inherent characteristic of the oscillatory system, which is determined by the given network configurations. Previous analytic studies discovered that a low (high) phase synchronization induces a large (small) phase response for a stimulus, proving that stimulation to the phases around a stable fixed point of the PRC increases phase response, whereas stimulation to the phases around an unstable fixed-point decreases phase response (*55*). These properties generally hold for networks with different coupling functions, network structures, and connectivity (*56*).

The present study advances previous results by demonstrating a role of fluctuations in network synchronization with the distance from criticality in the model, as well as empirical confirmation using EEG and fMRI from various states of consciousness, while directly calculating the amount of information sharing (*SMI*) and network susceptibility (*χ*) with the network synchronization *R* in the time domain. Here we proposed a role of the large synchronization fluctuation as a functional platform to create temporal windows processing internal or external information. In particular, the temporal windows characterized by large *SMI* and small *χ*, play a role in internal information integration, at the same time avoiding interventions from the outside world. These characteristics of a highly synchronized network create temporal windows shielded from external perturbations and simultaneously enable the brain to integrate globally distributed information across brain regions. Empirically, we demonstrated that such selective roles of temporal windows observed in the human brain networks in conscious states disrupt in altered states of consciousness. Notably, in our model study, we didn’t consider the neuromodulation system, which implies that the temporal windows for internal or external information processing can arise through the interplay between regional brain activities while the interactions are close to criticality. In other words, the emergence of the network preference for internal or external information and the continuous fluctuation between these two modes originate as a generic network feature near criticality, regardless of the specific neurobiology.

## Limitations

There are several limitations in this study. First, our brain network model simulated source signals of the brain network, not the EEG signals themselves in conscious and unconscious states that were tested. Instead, we focused on identifying the generic features of brain networks near and far from criticality to study the alteration of the brain network preference for internal/external modes in conscious and unconscious states. Second, we quantitatively evaluated the preferences for internal information and external stimuli without external stimulation. Since we cannot directly measure the response of unconscious subjects for cognitive tasks, which is beyond the scope of the current study, quantitative evaluations were limited to investigating the relationship between *R*, *SMI*, and *χ* at a system level. However, we expect that the results would be similar to the model results where we have already shown the relationship between R and responsiveness with external stimuli (*5*). Third, we focused on 4-12 Hz because this frequency band demonstrates global network features in conscious states that are suitable for the application of our brain network model (*25*). However, how the EEG of 4-12Hz is globally networked in the cortex and the relevant neuroanatomy is still unclear. Furthermore, higher frequency oscillations—which might be important for information processing—were excluded. Fourth, in our previous model study, we found more specific amplitudes and phases of oscillators that determine large and small responsiveness to external stimuli. However, in this study, we only used the level of synchronization as a criterion to classify the temporal windows due to the complexity of the application to the EEGs of various altered states of consciousness.

## Conclusion

Based on both a computational model and empirical data analyses from EEG and fMRI, we propose a novel role for criticality in temporal information processing in the brain. We found that a large synchronization fluctuation that spontaneously emerges near criticality enables to create temporal preferences for internal information and external stimuli. The distinct preferences at high and slow synchronizations were also found in EEG in the conscious state and our investigation of well-known fMRI networks supported the association of high and low synchronization periods with internal and external information processing in the brain. However, the temporal preferences vanish when the brain state deviates from criticality as well as in altered states of consciousness (ISO, KET, PSY, MCS, and UWS). The results lead us to conclude that the temporally fluctuating preferences create a functional platform for continuous switching between internal and external modes in the brain, which presumably plays an essential role in constructing inner models of the outside world in the conscious brain.

## Materials and Methods

### Simulation of spontaneous neural oscillations

A coupled Stuart-Landau model with human brain network structure has been widely used to simulate the oscillatory dynamics from various types of imaging modalities including EEG, MEG, and fMRI (*9*, *20*, *23*). Here we also used the coupled Stuart-Landau model to simulate the oscillatory dynamics of brain network and investigated relationships between network synchronization and variables associated brain network’s preferences for integrating information internally in the brain network and receiving stimuli from the outside of the brain.

Spontaneous networked oscillations were generated using a coupled Stuart-Landau model in a group-averaged anatomical human brain network constructed from diffusion tensor imaging (DTI) of 78 nodes (*57*). The coupled Stuart-Landau model, composed of the *N* number of oscillators, is defined as the following:

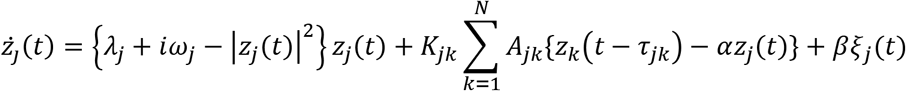

where a complex variable *z*_*j*_ (*t*) represents the oscillatory dynamics of brain region *j* at time *t*. *ω*_*j*_ is an initial angular natural frequency of oscillator j. We used Gaussian distribution for the natural frequency with a mean frequency of 10 Hz and a standard deviation of 0.5 Hz to simulate the peak frequency bandwidth of human EEG activity (*5*, *22*). *A*_*jk*_ = 1 if oscillators *j* and k are connected, and *A*_*jk*_ = 0 if they are not, based on the structural brain network. *τ*_*jk*_ is a time delay between oscillators *j* and *k*, defined as *D*_*jk*_/*s*, with the distance between brain regions *j* and *k*, *D*_*jk*_, and the average speed of axons in brain regions, *s*. Here we used *s* = 7*ms*. The node *j* receives input from connected node *k* after the time delay *τ*_*jk*_ . *λ*_*j*_ and *K*_*jk*_ are a bifurcation parameter of oscillator *j* and a coupling strength between oscillators *j* and *k*, respectively. Modulating these parameters induce competition of independent behavior of oscillator and the coupling among the oscillators so that complex oscillatory dynamics differently emerge in different parameter regions. Each node shows supercritical Hopf bifurcation and the dynamics of the oscillator settle on a limit cycle if *λ*_*j*_ > 0, and on a stable focus if *λ*_*j*_ < 0. We used a homogeneous bifurcation parameter *λ*_*jk*_ = *λ* from −3.2 to 3.2 with *δλ* = 0.2, and a coupling strength *K*_*jk*_ = *K* from 0 to 1 with *δK* = 0.02. We additionally modulated a diffusive coupling parameter *α*. The *α* controls the degree of outgoing flow of node *j*. In neural networks, two extreme values of *α*, 0 and 1 indicate two types of synapses, chemical synapses, and gap junctions, respectively. We tested 0, 0.5, and 1 for the *α*, and set *α* = 0.5 as the empirically well-fitted parameter. Further investigation for this parameter will be explored in the future study. *ξ*_*j*_(*t*) is a Gaussian white noise for each node *j* and added to the dynamics with the standard deviation *β* = 0.05. We numerically solved differential equations of the Stuart-Landau model using the Stratonovich-Heun method with 1,000 discretization steps, then we resampled the data with 500 Hz. The last 60 seconds were used for the analysis of each simulation after 15 seconds of saturation periods. Therefore, spontaneous oscillatory dynamics were generated for each brain region at each *λ* and each *K*. We repeated this simulation with one hundred different initial frequency distributions to obtain statistical stability.

### Experimental protocol and EEG acquisition

#### Isoflurane anesthesia

The isoflurane data were collected from a cohort of 20 healthy volunteers (20-40 yrs) recorded at the University of Michigan, Ann Arbor (Protocol #HUM0071578). The study has been reviewed by the Institutional Review Boards specializing in human subject research at University of Michigan. Written informed consent in accordance with the Declaration of Helsinki to participate in the study was obtained from all participants. Ten participants underwent general anesthesia. The participants in the anesthesia group initially received propofol at increasing infusion rates over three consecutive 5-min blocks (block 1: 100 μg/kg/min, block 2: 200 μg/kg/min, block 3: 300 μg/kg/min). Responsiveness was quantified every 30 s by the response to the verbal command “Squeeze your left/right hand twice” with random allocation to left/right. Isoflurane was then administrated with air and 40% oxygen at 1.3 age-adjusted minimum alveolar concentration. The isoflurane was administrated for 3 h and discontinued.

We analyzed EEG data of 9 out of 10 subjects who underwent general anesthesia due to the availability of high-density EEG data (128-channel HydroCel nets, Net Amps 400 amplifiers; Electrical Geodesic, Inc., USA). All EEG channels were referenced to a vertex with 500 Hz sampling frequency. EEG data of 4-min of eye-closed resting state before isoflurane administration (baseline) and 4-min of periods during general anesthesia (ISO) without burst-suppression were used in the current study. The data have been published with different analyses and hypotheses (*58*).

#### Ketamine anesthesia

The ketamine data were collected from 15 healthy volunteers (20-40 yrs) recorded at University of Michigan, Ann Arbor, with written informed consent to participate in the study. This study was approved by the University of Michigan Medical School Institutional Review Board, Ann Arbor, Michigan (HUM00061087). The data have been published with different analyses (*58*). EEG data were acquired during a 5-min eyes-closed resting state before ketamine administration, subanesthetic ketamine infusion (0.5 mg/kg) over 40 min, followed by 8 mg ondansetron, break for completion of questionnaire, anesthetic (1.5 mg/kg) bolus dose, and recovery period. EEG data of 4-min eyes-closed resting state (baseline), 4-min subanesthetic ketamine-induced state (PSY), and 4-min ketamine-induced unconsciousness (KET) after bolus anesthetic administration were used in the current study. The EEG data were acquired with 128-channel HydroCel nets, Net Amps 400 amplifiers (Electrical Geodesic Inc., USA). All channels were referenced to a vertex with 500 Hz sampling frequency.

#### Disorders of consciousness

EEG data were originally collected from a cohort of 80 patients with disorders of consciousness caused by ischemic stroke, intracerebral hemorrhage, subarachnoid hemorrhage, subdural hematoma, traumatic brain injury, meningitis, or hyperglycemic brain injury. Patients were diagnosed as minimally conscious or vegetative states/unresponsiveness wakefulness syndrome (UWS) using the Coma Recovery Scale-Revised (CRS-R). The CRS-R status was acquired again after six months of investigating the follow-up changes. The data from 17 subjects (the Munich cohort) were recorded on two different systems; 15 subjects were recorded with 64-channel, ring-type sintered, and nonmagnetic Ag/AgCl electrodes (Easycap, Herrsching, Germany); 2 subjects were recorded with 32-channel, nonmagnetic, and battery-operated electroencephalographic amplifiers (BrainAmp MR, Brain Products, Gilching, Germany). Both EEG data were recorded at 5 kHz sampling rates (BrainVision Recorder, Brain Products). The data from 63 patients (the Burgau cohort) were recorded with a 256-channel high-density Geodesic sensor net, a Net Amps 300 amplifier, and Net Station 4.5 software (Electrical Geodesic Inc., Eugene, OR, USA). The sampling rates were 1 kHz. All data were preprocessed to have 63-channel. In the current study, we analyzed the 4-min of the data from 9 subjects who were diagnosed as UWS without the evolution of CRS-R status and not showing suppression patterns and 16 subjects who were diagnosed as MCS.

### EEG data preprocessing

With three datasets, we analyzed 6 different states of consciousness, such as conscious state, (CS, n=24), ketamine-induced psychedelic state (PSY, n=15), isoflurane-induced unconsciousness (ISO, n=9), ketamine-induced unconsciousness (KET, n=15), MCS (n=16), and UWS (n=9). We selected 96-channel for the isoflurane and ketamine data and selected 56-channel for the MCS and UWS data that cover the scalp for the analysis. The MCS and UWS data were down-sampled to 250 Hz. After selecting the EEG channels, we high-pass filtered the data with 0.5 Hz cutoff frequency using MATLAB function “pop_eegfiltnew.m” in the EEGLAB toolbox. We then removed noisy epochs using MATLAB function “trimOutlier.m” with a standard deviation of channel amplitude rejection thresholds 100 μV, amplitude rejection thresholds 300 μV, and the range for rejection period 200-ms. Rejected channels were reconstructed using MATLAB function “pop_interp.m” in the EEGLAB with a spherical interpolation method. All EEG data from isoflurane, ketamine-induced unconsciousness, and disorders of consciousness were re-referenced to the average of all channels. We band-pass filtered the data with a frequency range from 4 to 12 Hz to capture the properties of globally coupled oscillators. We also analyzed the data with other frequency bands (1-4 Hz, 1-20 Hz, 8-12 Hz, 13-30 Hz, and 30-42 Hz).

### Level of network synchronization and pair correlation function

In this study, the level of network synchronization is an important factor to determine the brain network’s preference for processing internal information of the network or external information from the outside of the brain network. The instantaneous level of network synchronization *r*(*t*) at time *t* was measured by the order parameter,

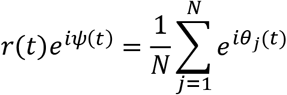

where *θ*_*j*_(*t*) is a phase of oscillator *j* and *φ*(*t*) is the average global phase at time *t*. Here *r*(*t*) equals to 0 if phases of oscillators are uniformly distributed and 1 if all oscillators have the same phase. The *r*(*t*) was calculated for all parameter combinations in the model, all states from EEG, and fMRI data.

We then measured a pair correlation function *PCF* ≡ *N*[< *r*^2^(*t*) > −< *r*(*t*) >^2^], which is a surrogate measure of criticality, to define the critical state(*18*) in the model, under the assumption that the largest synchronization fluctuations are associated with the largest number of metastable states of brain network and should be shown at criticality. We measured PCF for all parameter combinations in the model, all states of EEG data.

### Temporal window classification based on the network synchronization

To investigate the network’s information processing preference with reliable time resolution, we classified the network transient states into high and low *R* temporal windows. We calculated an average network synchronization *R* as < *r*(*t*) >_*T*_ for each temporal window, where *T* is the size of the temporal window. In the model, we set *T* = 250 *msec* with 50 *msec* overlaps and classified the temporal windows into two different windows: one of which is the incoherent state and the other is the highly synchronized state (low and high *R* windows). In the model, we set thresholds as *R* = 0.3 and *R* = 0.5 for low and high*R* windows, respectively. For the EEG data, we set *T* = 250 *msec* with 50 *msec* overlaps for CS, PSY, ISO, and KET, and *T* = 300 *msec* with 50 *msec* overlaps for MCS and UWS due to the sampling frequency of the data. We concatenated *R* values across all temporal windows, all subjects, and all six different states of consciousness to find the total mean and standard deviation of *R*. With thresholds 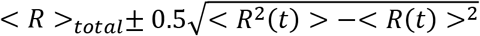, we classified all temporal windows across all states into windows with low and high *R*.

### Symbolic mutual information and Susceptibility

To calculate the information processing capabilities for each temporal window, we measured symbolic mutual information (SMI) and network susceptibility in the model and EEG signals. The SMI measures the amount of information sharing, quantifying the extent of non-random joint fluctuations between two signals *X* and *Y*. To calculate the SMI, the signals *X* and *Y* are first transformed into sequences of discrete symbols 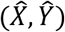 with a fixed symbol size *m* with a temporal separation *τ*. It calculates a joint probability of co-occurring symbolic patterns between two signals.

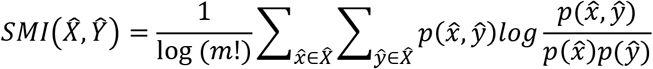

where 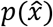 is the probability occurring symbol 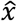 in the time series 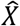, leading to We set *m* = 3, leading to 3! = 6 of different symbol pairs 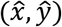 that can potentially exist in the transformed symbolic time-series. In the model, we used *τ* = 14 (28 *ms*) to get the maximum resolved frequency 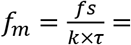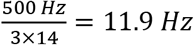, which is suitable for our interest of frequency range (~12 Hz) (*59*). The SMI was calculated between all node pairs in the model (all channel pairs of the EEG signals) for all temporal windows we defined above. We took an average across SMI values between pairs and defined the average SMI as the amount of total information sharing in each temporal window. For the EEG signals, we compared pairwise SMI values of real data to the pairwise SMI values of twenty surrogate data set with phase randomization technique and used only statistically significant SMI values (p<0.01, Wilcoxon rank-sum test). The average SMI value of EEG channel pairs was used as an amount of total information sharing in each temporal window.

The network susceptibility was also measured in both model and EEG signals. The network susceptibility *χ* was defined as following (*18*):

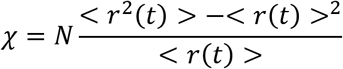

We defined the stationary dynamical susceptibility *χ* for each temporal window to see how susceptible the brain state would be to the external information from the outside of the brain in each temporal window.

### Topographic similarity

To understand distinct functional network configurations engaged in low and high *R* windows, we measured topographic similarity *ρ*^*amp*^, which is defined as Spearman correlation between node degrees and node amplitudes.

In the model, the degree of brain regions was defined as the number of structural connections between one region and the other regions. The instantaneous amplitude *Z*(*t*) was measured by the absolute value of the complex variable *z*(*t*). Th e in stantaneous Sp earman co rrelation between degree and amplitude, *ρ*^*amp*^(*t*), was calculated for each time point for 60 seconds for all parameter combinations of the model. Then we took average of *ρ*^*amp*^(*t*) for e ach tempo ral window. For the EEG data, the degree of a node (channel) was inferred from the average functional connectivity strength over time measured by weighted phase lag index (wPLI) within a frequency range from 4 to 12 Hz for each subject. The wPLI is a measure that captures phase locking between two signals *X* and *Y*, relatively robust to volume conduction of EEG (*60*).

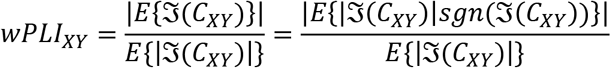

where 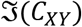 is an imaginary part of cross-spectrum *C*_*XY*_ between two signals *X* and *Y*. Here we used Hilbert-transformed complex signals for the calculation of cross-spectrum *C*_*xy*_. The *wPLI*_*XY*_ equals to 1 if the phases of one signal *X* always lead or lag the phases of the signal *Y*, and equals to 0 if the phase lead/lag relationship between two signals is perfectly random. We constructed the wPLI matrix across all channels for each 30-second epoch with a 5-second overlap and binarized the wPLI matrix for each epoch with the top 30 % wPLI values. We calculated the degree of channels for each epoch and took the average over all epochs so that we can capture the statistically intuitive structural degree that reflects the strength of neural communication across brain regions associated with each EEG electrode. Note that the degree is extracted from the CS (baseline) for each subject and applied to the analysis for the altered states of consciousness induced by isoflurane and ketamine. Since MCS and UWS patients have no baseline, we excluded those data set for this analysis. The instantaneous amplitude was calculated by the absolute value of the Hilbert transformed EEG signals from a frequency range of 4 to 12 Hz. The instantaneous Spearman correlation between degree and amplitude *ρ*^*amp*^(*t*) was calculated for each subject and CS, ISO, and KET.

### Correlation between network synchronization, symbolic mutual information, network susceptibility, and topographic similarity

According to our hypothesis, levels of network synchronization are associated with the network’s preference for internal and external modes in the brain on a sub-second time scale. Therefore, we calculated Spearman correlation between the level of network synchronization and the information processing metrics (SMI and *χ*). We also calculated Spearman correlation between the level of network synchronization and the topographic similarity. The correlation between *R*, *SMI*, *χ*, and *ρ*^*amp*^ were calculated across all temporal windows. For the model, we calculated Spearman correlations for each parameter combinations with one hundred different simulations (Fig. 2). For the EEG data, we calculated Spearman correlations between *R*, *SMI*, *χ*, and *ρ*^*amp*^ for each subject and each state to investigate whether the relationships we found from the model hold for the empirical EEG data (Fig. 5).

### Joint histogram between SMI and *χ*

To visualize the distinct network preferences for internal and external processing modes, a joint histogram of *SMI* and *χ* of low and high *R* windows was calculated. The joint histogram for the model was calculated near and far from the critical state across all temporal windows of one hundred frequency distributions with the bin size 0.02 for *SMI* and 0.1 for *χ* (Fig. 3e). For the EEG data, we calculated a joint histogram of SMI and *χ* for each temporal window with the bin size 0.02 for SMI and 0.1 for *χ* during CS (Fig. 4f), ISO, KET, PSY, MCS, and UWS (Fig. 5f). We can easily figure out the temporal window’s preference for internal or external information by calculating the joint histogram.

### Power-law analysis for dwell-time of the positive correlation between degree and amplitude

It has been known that one of the characteristic features of criticality is the power-law distributions of dynamics. The dynamics following probability distribution *p*(*x*) ∝ *x*^−*β*^ imply that all values *x* can occur without a characteristic size or scale (*61*). It has been also known that the frequency density of phase-locking intervals and the change in the number of phase-locked pairs between successive time points display power-law distributions at criticality in the Ising model and Kuramoto model (*6*). Since the network synchronization is correlated with the topographic similarity *ρ*^*amp*^ (relationship between functional network configuration constrained by network structure in the brain), we estimated the probability distribution of dwell-time of *ρ*^*amp*^ to check whether the relationship between functional network configuration and network structure follows power-law at during conscious wakefulness. For the EEG data, we measured dwell-time of positive correlation periods (hub-dominant configuration periods) across all subjects for each state (Fig. S9). We used a python package “powerlaw” (https://pypi.org/project/powerlaw/) to estimate the probability distributions of positive correlation periods. We fitted dwell-time distributions to power-law and exponential distributions and compared their loglikelihood values to determine which distribution is well-fitted to the data. Using the loglikelihood ratio R_*L*_ and p-value, we obtained the estimated probability distribution with the exponent *β*. R_*L*_ has a positive value if the power-law fit is more appropriate, while it has a negative value if the exponential fit *p*(*x*) ∝ *e*^−*β*^ is more appropriate. We set the minimal value *x*_*min*_ as 200-ms for the EEG data to match the speed of the brain network dynamics and focused on a “heavy-tailed” characteristic of power-law.

### fMRI experimental protocol, data acquisition, and preprocessing

The fMRI data were collected at two different research sites, Wisconsin and Shanghai. The experimental protocol for the first data set recorded from Wisconsin was approved by the Institutional Review Board of Medical College of Wisconsin (MCW). Propofol was administrated to 15 healthy volunteers (male/female 9/6; 19-35 yrs) and the OAAS (observer’s assessment of alertness/sedation) was quantified to measure the level of behavioral responsiveness. This dataset has been previously analyzed with a different hypothesis and published (*19*). In the current study, we used conscious states (baseline) that subjects responded readily to verbal commands (OAAS score = 5) and deep sedation that subjects have no response to verbal commands (OAAS score = 2-1). At the deep sedation level, the plasma concentration of propofol was maintained at equilibrium by continuously adjusting the infusion rate. The corresponding target plasma concentrations of propofol vary across subjects (1.88±0.24 μg/ml) due to the individual variability of anesthetic sensitivity. Total 14 subjects were analyzed in the current study because one subject had to be excluded due to excessive movements. Rs-fMRI data of the conscious state and deep sedation both consisted of a 15-min scan. Gradient-echo EPI images of the whole brain were acquired on a 3T Signa GE 750 scanner (GE Healthcare, Waukesha, Wisconsin, USA) with a standard 32-channel transmit/receive head coil (41 slices, TR/TE = 2000/25 ms, slice thickness = 3.5 mm, field of view = 224 mm, flip angle = 77°, image matrix: 64×64). Rs-fMRI was co-registered by high-resolution anatomical images. See (*19*) for a more detailed experimental protocol.

The second dataset includes 28 healthy participants (male/female 14/14) and 21 patients with disorders of consciousness (male/female 18/3). The study was approved by the Institutional Review Board of Huashan Hospital, Fudan University. The healthy controls had no history of neurological or psychiatric disorders or were taking any kind of medication. The patients were diagnosed as either minimally conscious or in the vegetative state/unresponsive wakefulness syndrome (UWS) according to the Coma Recovery Scale-Revised (CRS-R) on the day of fMRI scanning. We analyzed the data of 13 patients who were diagnosed as UWS in the current study. This dataset also has been published using different a hypothesis (*19*). Gradient echo EPI images of the whole brain for the second dataset were acquired on a Siemens 3T scanner (Siemens MAGNETOM, Germany) with a standard 8-channel head coil (33 slices, TR/TE = 2000/35 ms, slice thickness = 4 mm, field of view = 256 mm, flip angle = 90°, image matrix: 64×64). Total two hundred EPI volumes (6 minutes and 40 seconds) were acquired with high-resolution anatomical images.

Preprocessing steps were implemented in AFNI (http://afni.nimh.nih.gov/), which included slice timing correction, head motion correction/realignment, frame-wise displacement estimation, coregistration with high-resolution anatomical images, spatial normalization into Talaraich stereotactic space, high-pass filtering (>0.008 Hz), regressing out undesired components (e.g., physiological estimates, motion parameters), spatial smoothing (6 mm full-width at half-maximum isotropic Gaussian kernel), temporal normalization (zero mean and unit variance). Global signal regression (GSR) was not applied to preserve whole-brain patterns of co-activation or co-deactivation. More detailed preprocessing steps were described in the paper (*19*).

### Co-activation pattern analysis for fMRI data

An unsupervised machine-learning approach named k-means clustering algorithm was used to define co-activation patterns (CAPs). The algorithm operates as a classifier of a set of objects (e.g. fMRI volumes) to minimize within category (e.g. patterns) differences and maximize across category differences. Specifically, the fMRI volumes were pooled together across states (baseline, propofol-induced states, ketamine-induced states, and UWS) and subjects, and classified into k number of clusters (patterns) based on their spatial similarity, yielding a set of CAPs or brain states (*19*). As such, we obtained time-series of discrete CAP labels from the original fMRI time-series (voxels×volumes). The volumes containing motion artifact tagged by motion censoring procedure were not included in the CAP analysis. We evaluated the validity of the number of clusters by measuring the inter-dataset similarity (Euclidean distance) of the average CAP occurrence rate distributions of conscious and unresponsive conditions. We searched the number of k from 2 to 30 using an index, (CC + UU)/(2 × CU), as the ratio of inter-dataset similarity among conscious conditions (CC) and among unresponsive conditions (UU) versus the inter-dataset similarity among conscious and unresponsive conditions (CU) across all datasets and found the optimized number of k as 8. Therefore, each fMRI volume from all datasets corresponded to one of the patterns from 8 CAPs. In the current study, we used pre-defined CAPs and analyzed a subset (n=69) of originally published datasets including subjects under wakefulness baseline condition (n=42), deep sedation induced by propofol (n=14), and UWS patients (n=13).

### Dominant CAPs for incoherent and highly synchronized states

In the EEG data, we defined the incoherent and highly synchronized state that presumably corresponds to externally susceptible/internally integrated state using certain thresholds of the level of synchronization. To define the incoherent/highly synchronized state for the fMRI data, we also measured the level of synchronization r(t) across voxels for each fMRI volume. Then we classified fMRI transient state to incoherent or highly synchronized state using the thresholds 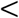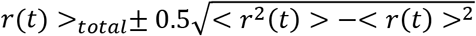. The < *r*(*t*) >_*total*_ were calculated from all concatenated fMRI volumes of three different states of consciousness (baseline, propofol-induced deep sedation, and UWS). Then we investigated occurrence rate distributions of CAPs for the incoherent or highly synchronized state with a permutation test. For the permutation test, we generated null CAP time-series by 20000 permutations, randomly and uniformly exchanging CAPs in time Then the original fMRI time points (volumes) that levels of synchronization have below (above) threshold 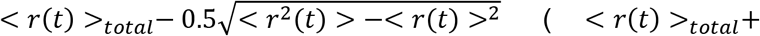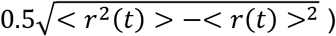for each state of consciousness were selected and patterns corresponding to those time points were used to examine whether the occurrence probabilities significantly deviate from uniformly random sequences. The significantly dominant CAPs were determined at the significance level of p<0.005 (0.5% percentile of the null distributions: one-sided).

## Acknowledgement

This research was supported by the National Institute of General Medical Sciences of the National Institutes of Health (Bethesda, MD) under award numbers R01GM111293 (ketamine data) and R01GM098578 (computational model), and the James S. McDonnell Foundation, St. Louis (isoflurane data), and by the Department of Anesthesiology, University of Michigan, Ann Arbor, Michigan. The authors also thank the Klinikum rechts der Isar, Technische Universität München (Munich, Germany) and the Therapiezentrum Burgau (Burgau, Germany) for providing the MCS and UWS EEG data.

