## Supplemental Figure 1-9 for "Criticality Creates a Functional Platform for Network Transitions between Internal and External Processing Modes in the Human Brain"

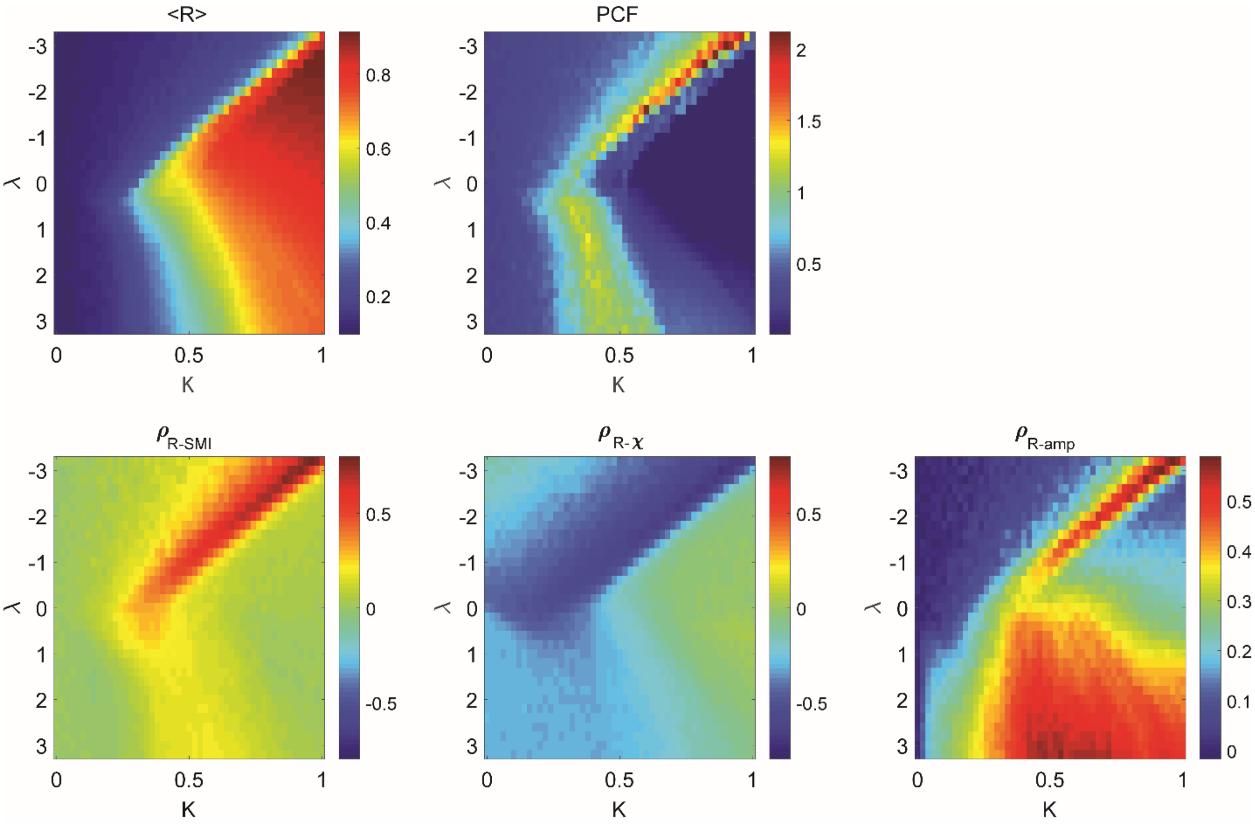


**Figure S1**. A level of global network synchronization *R* over 60 seconds, pair correlation function (PCF), and Spearman correlation between *R* and *SMI*, *R* and $\chi$, and *R* and $\rho^{amp}$ of the brain network model with the diffusive coupling strength $\alpha = 0.5$. The critical states are defined as the states with the maximum PCF. The brain network model shows maximum positive (negative) correlations between *R* and *SMI* (*R* and $\chi$) near critical states. The topographic similarity $\rho^{amp}$ shows maximum positive correlations with *R* near the critical states. The results were calculated from the average of 100 different initial frequency configurations.

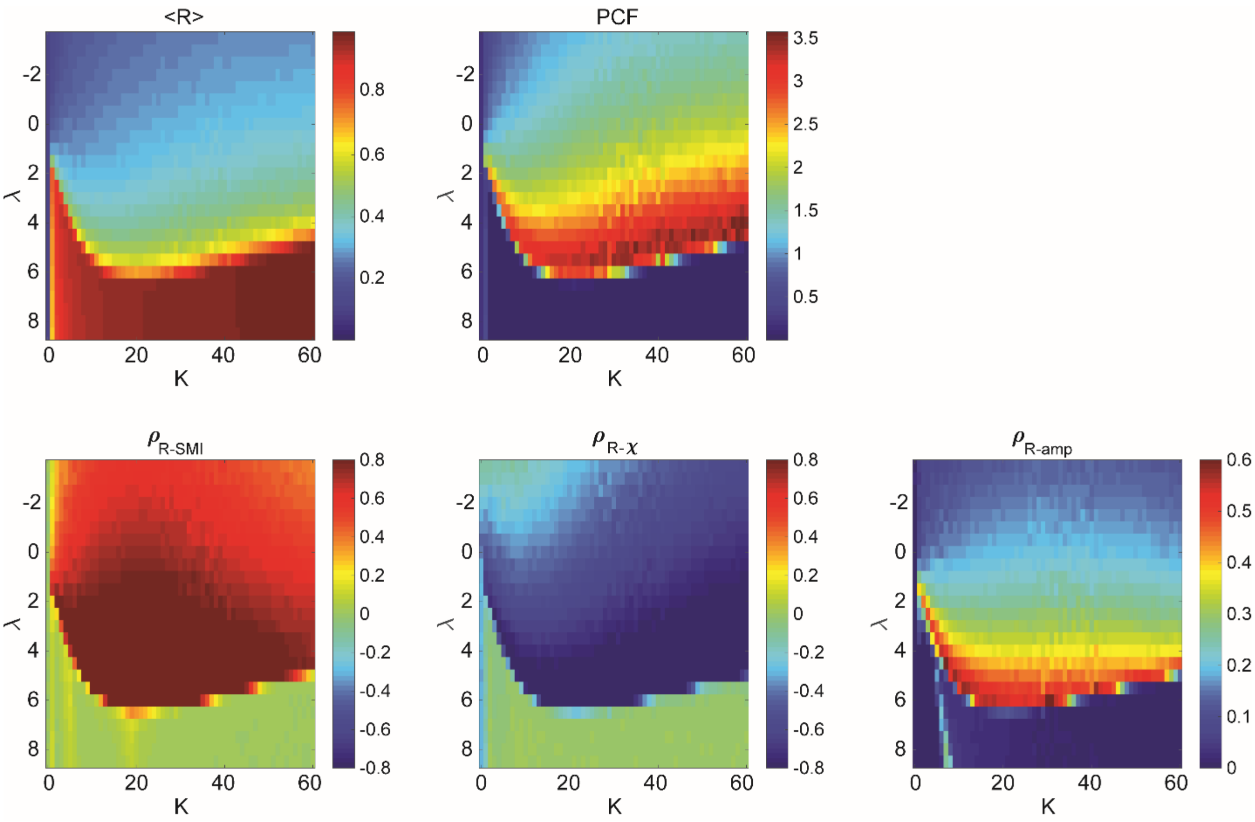


**Figure S2**. A level of global network synchronization *R* over 60 seconds, pair correlation function (PCF), and Spearman correlation between *R* and *SMI*, *R* and $\chi$, and *R* and $\rho^{amp}$ of the brain network model with the diffusive coupling strength $\alpha = 1$. (coupled in perfectly diffusive way). The critical states are defined as the states with the maximum PCF. The brain network model shows maximum positive (negative) correlations between *R* and *SMI* (*R* and $\chi$) near critical states. The topographic similarity $\rho^{amp}$ shows maximum positive correlations with *R* near the critical states. The results were calculated from the average of 100 different initial frequency configurations.

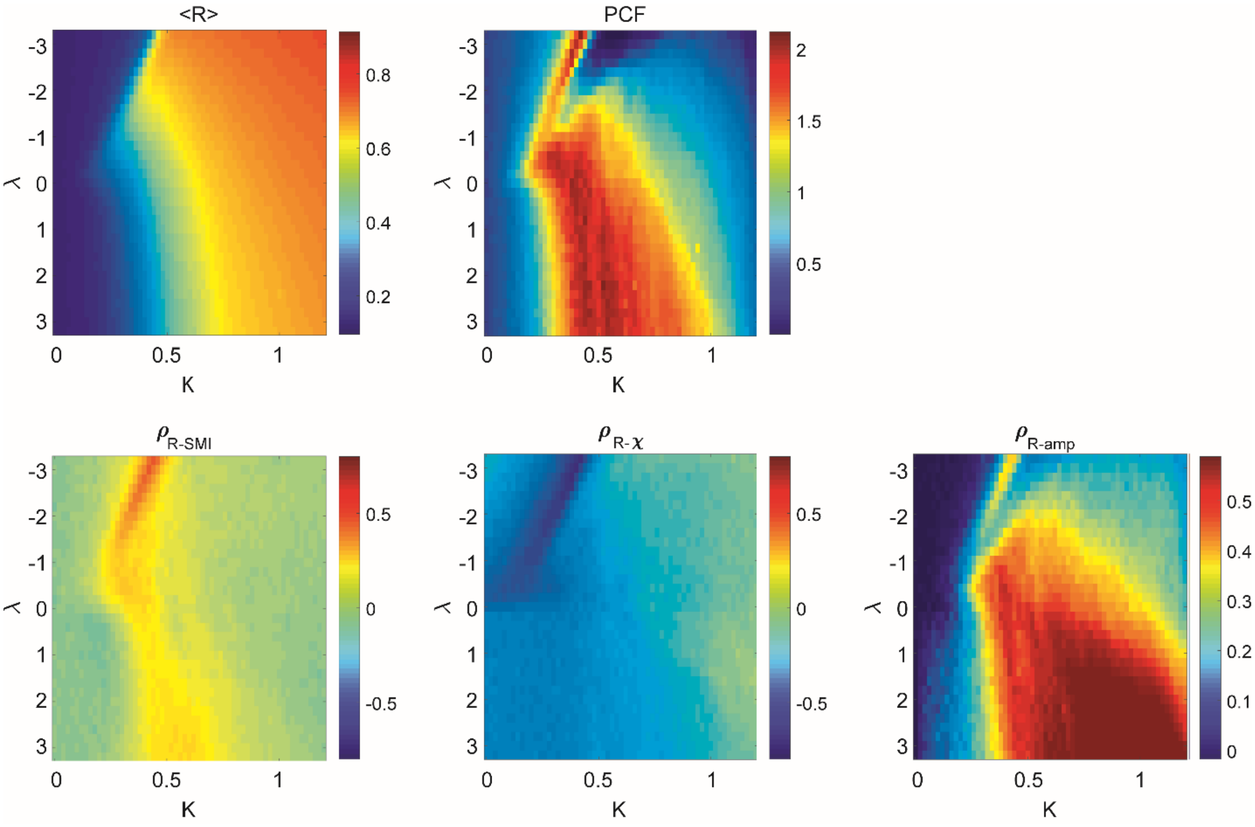


**Figure S3**. A level of global network synchronization *R* over 60 seconds, pair correlation function (PCF), and Spearman correlation between *R* and *SMI*, *R* and $\chi$, and *R* and $\rho^{amp}$ of the brain network model with the diffusive coupling strength $\alpha = 0.$ (coupled in perfectly direct way). The critical states are defined as the states with the maximum PCF. The brain network model shows maximum positive (negative) correlations between *R* and *SMI* (*R* and $\chi$) near critical states. The topographic similarity $\rho^{amp}$ shows maximum positive correlations with *R* near the critical states. The results were calculated from the average of 100 different initial frequency configurations.

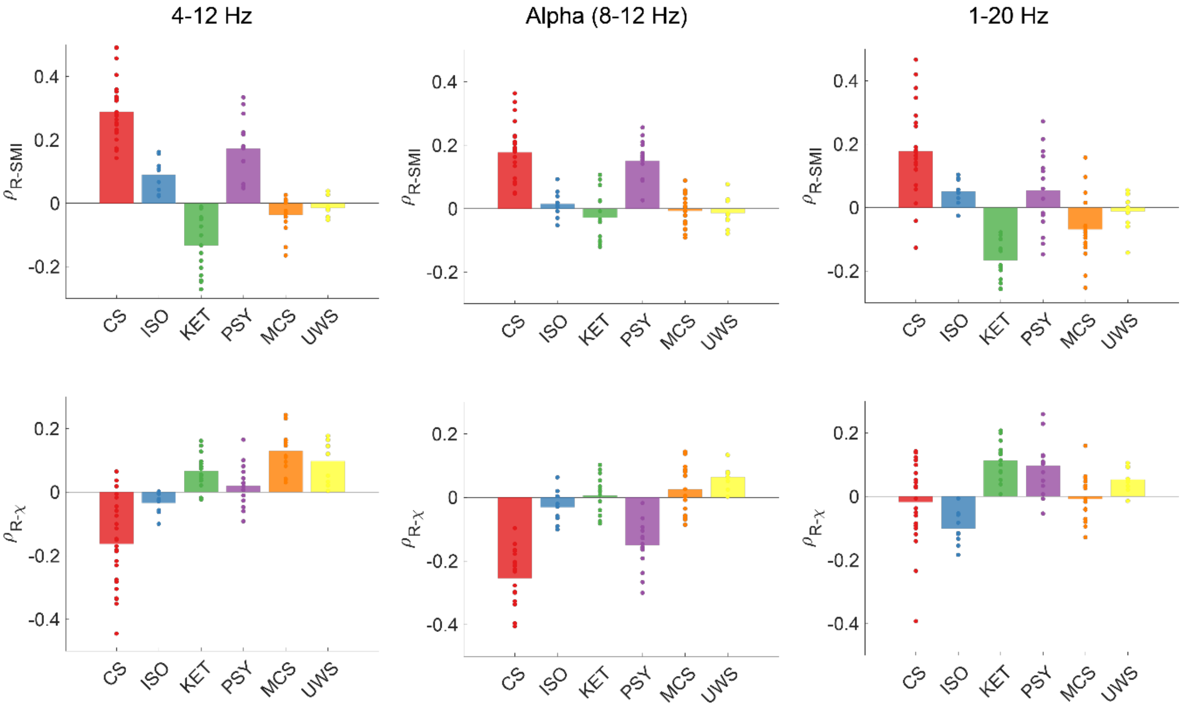


**Figure S4**. Spearman correlations between *R* and *SMI* (upper panels) and $R$ and $\chi$ (lower panels) in different states of consciousness with different frequency ranges. Dots indicate the correlation values of subjects. The relationships we found from the frequency range of 4-12Hz are maintained in alpha frequency range (8-12 Hz), but not in the frequency range 1-20 Hz. It suggests that the relationships between global network synchronization and brain network's preferences for internal and external information processing is a general property of globally networked oscillators.

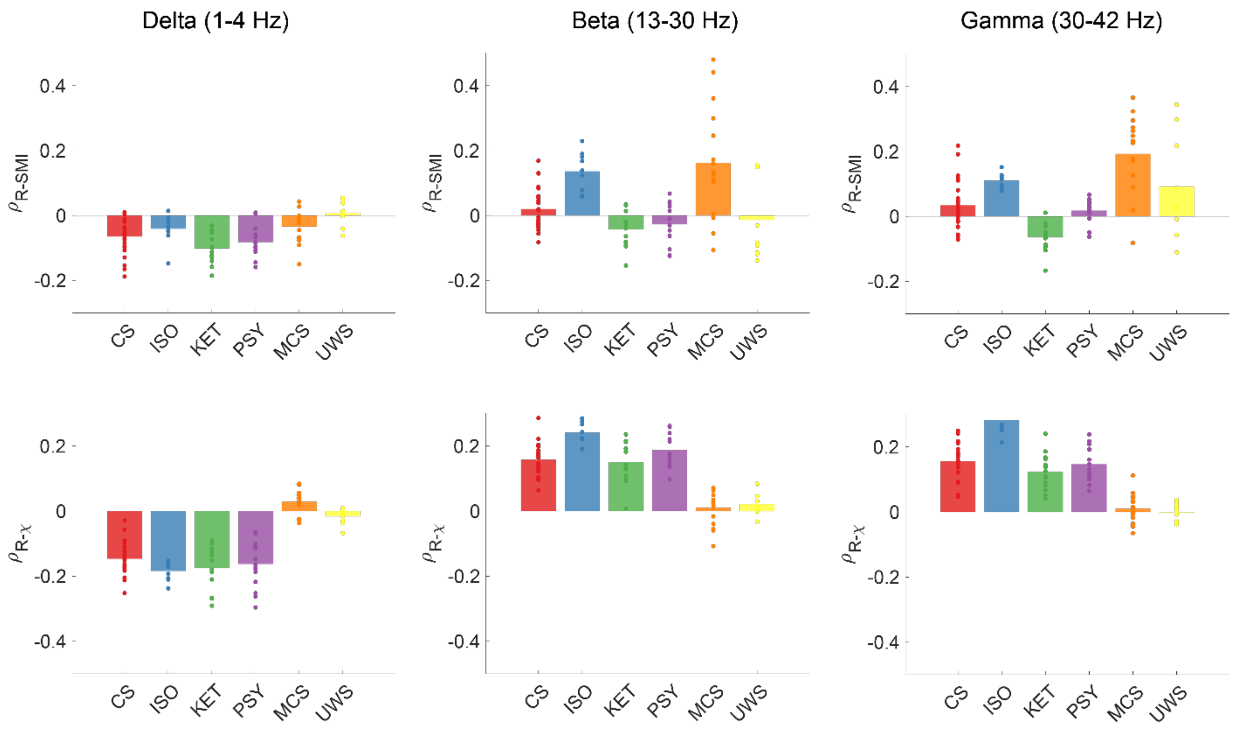


**Figure S5**. Spearman correlations between *R* and *SMI* (upper panels) and *R* and $\chi$ (lower panels) in different states of consciousness with different frequency ranges. Dots indicate the correlation values of subjects. The relationships we found from the frequency range of 4-12Hz are not maintained in the frequency range 1-4 Hz (delta), 13-30 Hz (beta), and gamma (30-42 Hz). It suggests that the relationships between global network synchronization and brain network's preferences for internal and external information processing is a general property of globally networked oscillators.

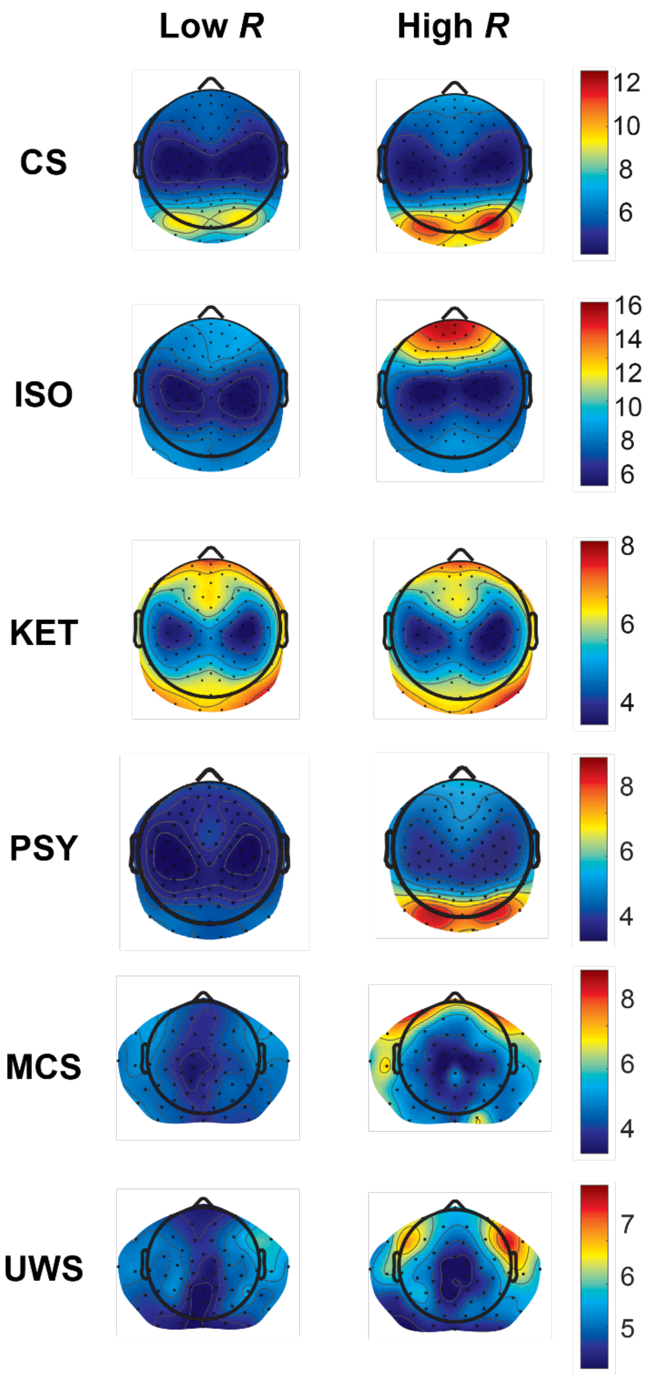


**Figure S6**. Functional brain network configurations of temporal windows of low and high $R$ in different states of consciousness. During CS (baseline), the amplitudes of posterior area (hub regions) in high $R$ windows are larger than the amplitudes in low $R$ windows, with the positive $\rho^{amp}$ for high $R$ windows.

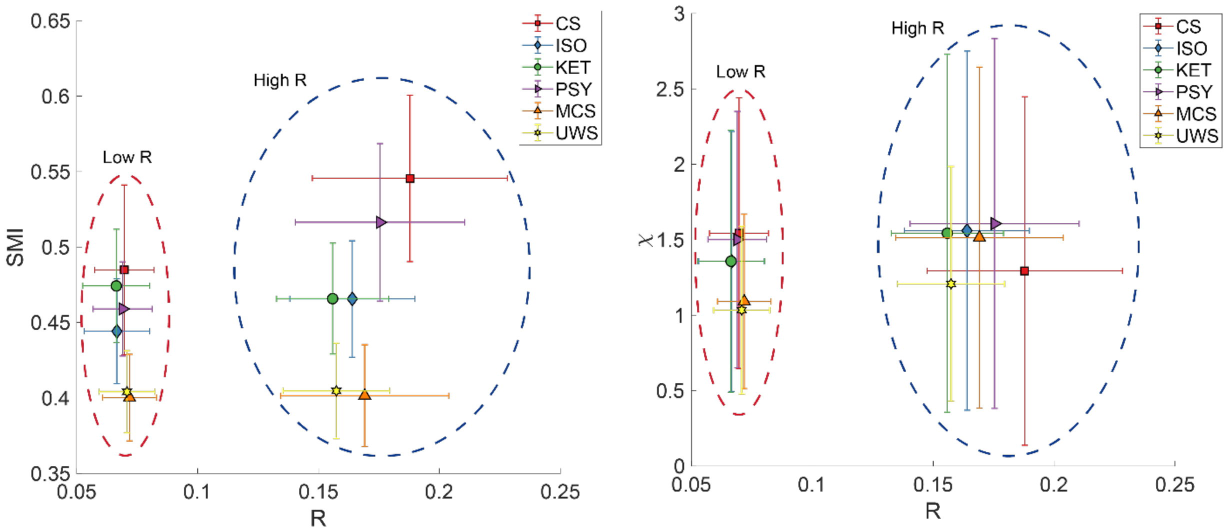


**Figure S7**. Comparisons of *R*, *SMI*, and $\chi$ values across different states of consciousness. Red (blue) dashed circles indicate temporal windows of low *R* (high *R*). Error bars indicate SD. The *R* values across all states are similar to each other for low and high *R* windows, respectively. The *SMI* values in high *R* window are different to each other, suggesting the amount of shared information in the brain network is different across the states even with the similar levels of *R*, suggesting the functional network configurations of the brain network are also important for internal information integration.

**CAPs (co-activation patterns)**

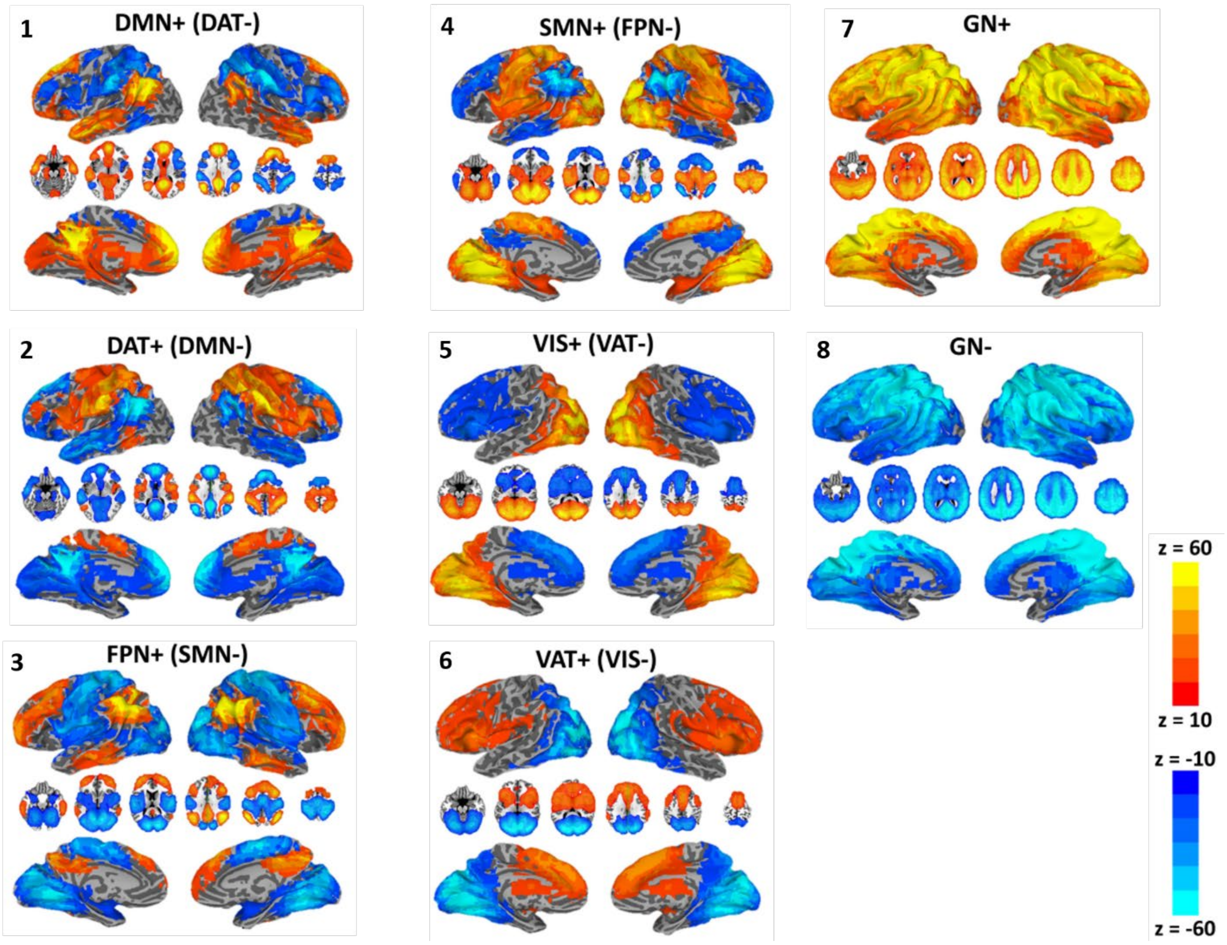


**Figure S8**. Spatial maps of fMRI co-activation patterns (CAPs). The CAPs consist of eight different functional networks such as a default-mode network (DMN+), dorsal attention network (DAT+), frontoparietal network (FPN+), sensory and motor network (SMN+), visual network (VIS+), ventral attention network (VAT+), and global network of activation and deactivation (GN+ and GN-). The figure is adapted with permission from the reference (60) Z. Huang, J. Zhang, J. Wu, G. A. Mashour, A. G. Hudetz, Temporal circuit of macroscale dynamic brain activity supports human consciousness. *Sci. Adv.* **6**, 87–98 (2020).

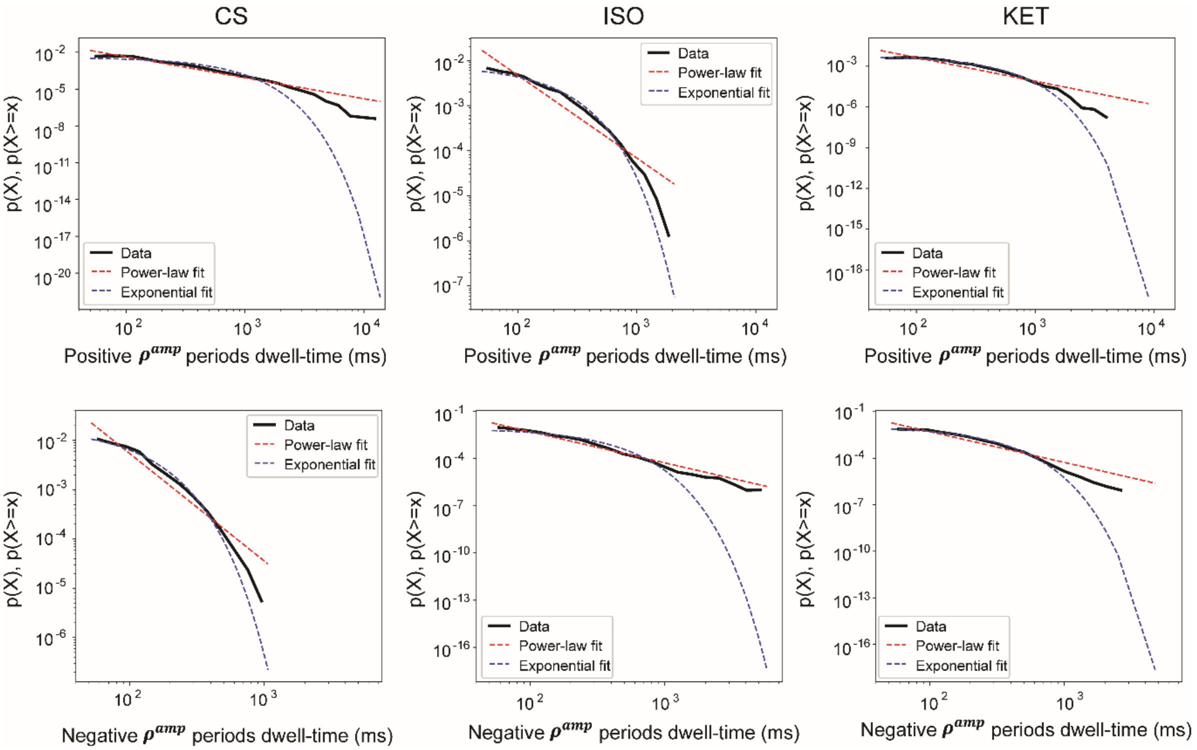


**Figure S9**. Probability density functions of $\rho^{amp}$ dwell-time and fitted power-law and exponential distributions in CS, ISO, and KET. A dwell-time of positive (negative) $\rho^{amp}$ periods was calculated across all subjects. The upper panel shows positive $\rho^{amp}$ dwell-time in CS (left), ISO (middle), and KET (right). Black lines indicate the data, red (blue) lines indicate power-law (exponential) distribution-fitted lines. The positive $\rho^{amp}$ in CS follows power-law, while that in ISO and KET is more likely to follow exponential distribution. The lower panel shows negative $\rho^{amp}$ dwell-time in CS, ISO, and KET.
